# The Preoptic Area and Dorsal Habenula Jointly Support Homeostatic Navigation in Larval Zebrafish

**DOI:** 10.1101/2023.05.18.541289

**Authors:** Virginia Palieri, Emanuele Paoli, Ilona C Grunwald Kadow, Ruben Portugues

**Author notes:** co-corresponding authors Correspondence to: Ruben Portugues, Ilona Grunwald Kadow. equal contribution (Virginia Palieri, Emanuele Paoli).

## Abstract

Animals must maintain physiological processes within an optimal temperature range despite changes in their environment. While the preoptic area of the hypothalamus (PoA) acts as a thermostat in mammals through autonomic and behavioral adaptations, its role in temperature regulation of animals lacking internal homeostatic mechanisms is not known. Through novel behavioral assays, wholebrain functional imaging and neural ablations, we show that larval zebrafish achieve thermoregulation through movement and a neural network connecting the PoA to brain areas enabling spatial navigation. PoA drives reorientation when thermal conditions are worsening and conveys this information for instructing future motor actions to the navigation-controlling habenula (Hb) - interpeduncular nucleus (IPN) circuit. These results suggest a conserved function of the PoA in thermoregulation acting through species-specific neural networks. We propose that homeostatic navigation arose from an ancient chemotaxis navigation circuit that was subsequently extended to serve in other sensory modalities.

## Introduction

Movement with the goal to relocate in response to environmental perturbations is used by most animals to maintain their physiological processes within a healthy range (i.e. homeostasis)^1^. Regardless of the specific strategy and environmental factors involved, the nervous system plays a crucial role in preserving homeostasis by comparing environmental and internal signals relative to past and ongoing experiences and physiological states^2^. Temperature is a prototypical example of how an environmental factor can critically affect the physiological processes of all types of organisms^3^. A failure to assess its value, valence, and rate of change can have a variety of consequences, from tissue damage to failure of the entire system^3–5^. Endotherms, such as mammals, can couple volitional behavioral strategies, for example warm or cold-seeking behavior, with autonomic measures, such as shivering/sweating or vasoconstriction/vasodilation^6^, to cope with thermal stress. On the contrary, ectotherms, such as fish, amphibians and reptiles, lack on autonomic homeostatic mechanism to regulate their body temperature^7^. These animals need to move in order to navigate towards places where the external temperature matches their homeostatic setpoint^8^. Thus, for ectotherms, thermoregulation is achieved by and large through navigation. We will refer to this phenomenon as homeostatic navigation.

Despite extensive research into the behavioral strategies and brain regions involved in either navigation or homeostasis, the manner in which these two processes interact and how the nervous system supports homeostatic navigation is poorly understood. Notably, it is an open question whether brain areas such as the preoptic area of the hypothalamus (PoA) regulating temperature homeostasis in endotherms also regulate homeostatic navigation in ectothermic organisms.

In this study, we aimed at dissecting the characteristics and neural mechanisms of homeostatic navigation in larval zebrafish using a combination of spatial, temporal and virtual thermal gradients. We show that fish modulate their reorientation probability and direction of turning based on their sensory context and previous motor choices. In this way, when the context is worsening (i.e. going away from the temperature setpoint), the reorientation probability significantly increases and fish favor coherent, steering reversal maneuvers over random turns or immobility.

Using a combination of whole-brain calcium imaging, circuit perturbations and behavioral phenotyping we identified two evolutionary conserved brain structures, the PoA and the Dorsal Habenula (dHb, a homolog of mammalian medial Hb), as critical players for homeostatic navigation. Fish with an ablated PoA showed a reduced reorientation probability upon worsening temperature and showed a strong impairment in coherent directional swims (U-maneuvers) observed in wild type and controls.

By contrast, dHb-ablated fish retained the drive to reorient when sensory conditions were worsening but showed the same impairment to move in a coherent direction away from unfavorable temperatures as observed in the PoA ablation experiments. Furthermore, in line with a modality-independent role in homeostatic navigation, we find that neurons in a small region of the right dHb respond to both thermal and salinity changes with similar dynamics. Thus, these neurons might convey multimodal input to downstream structures.

Based on our experimental data, we propose a model where the PoA and dHb are both involved in homeostatic navigation and support larval zebrafish thermoregulation. The PoA promotes an acute, sensory-based and directional increase in reorientation probability when environmental temperature moves away from its setpoint. This information is then passed to the dHb, which couples the memory of the sensory experience to previous motor choices to drive coherent movement trajectories away or toward the preferred temperature. Moreover, the dHb-IPN pathway appears to be a generalist circuit conveying a more abstract representation of the stimulus context independent of modality.

## Results

### Zebrafish perform homeostatic navigation by modulating the reorientation probability and direction

We first established that larval zebrafish control their body temperature by navigating a shallow temperature gradient without the need for other sensory information. To this end, we designed a 20 x 4 cm rectangular arena (Supplementary figure 1a) where we selectively heated up a 4 x 4 cm area at one end to 33 °C, effectively establishing a linear thermal gradient (see Methods and Supplementary figure 1b) of 0.04 °C/mm along the long side of the arena (Supplementary figure 1c). In this environment, the temperature difference across the length of the fish’s body would be maximal 0.2 °C. We monitored individual 5 - 7 days post fertilization (dpf) zebrafish for 15 minutes (see Methods) and found that by the end of the experiment, fish robustly avoided the hot side of the arena (Figure 1a, top and bottom panel). We then extracted the temperature experienced by the fish from its position in the arena (see Methods). We observed that, after an initial exploratory phase of approximately 260 s, fish rapidly appear to prefer a temperature (T_pref_) of ∼25.3 °C which we posit to be their homeostatic setpoint (Figure 1b and Supplementary figure 1d, f, g, and e for single example fish trajectories).

**Figure 1:**
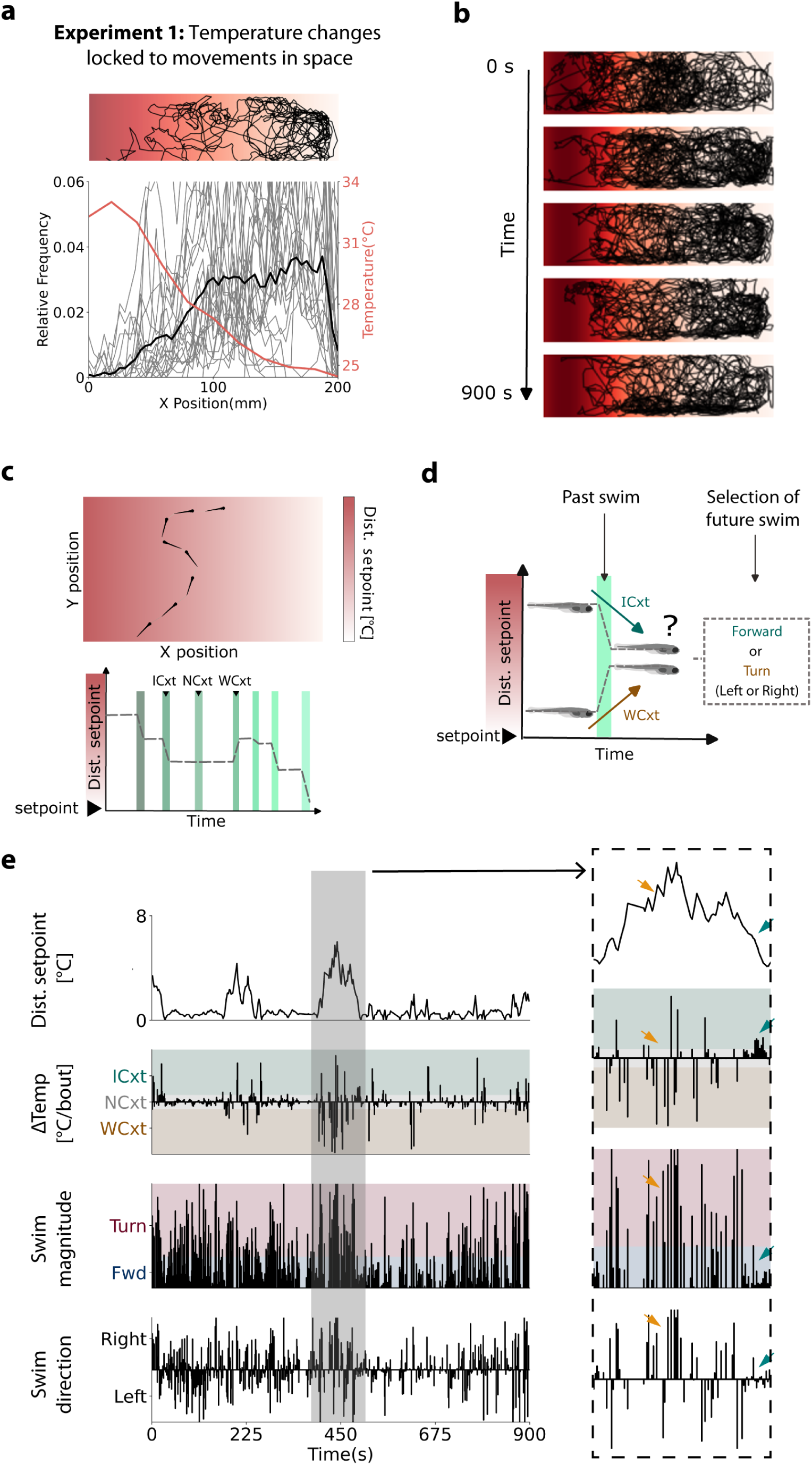
Larval zebrafish maintain homeostasis in a linear shallow thermal gradient. **a**. Top: example trajectories of 3 fish for the entire duration of the experiment (15 min). Actual temperature measured in the water is pictured with different shades of red. The highest temperature (33°C) is represented on the left by the dark red. Bottom: Histogram of animals’ x-position in the arena (n=26, gray traces), median of the population (black trace) and temperature measurement (red trace). **b.** Trajectories of all the fish split in 3-minute bins. **c.** Top: Sketch of an example trajectory. Bottom: change of temperature perceived over time for each swim event in the top sketch. Time is coded by the shades of green (from dark to light green). Arrows on top point to a movement in a ICxt (improving context), NCxt (no context) and a WCxt (worsening context), respectively. **d.** Sketch of information available to the fish for behavioral choice using the sensory context provided by the previous swim. **e.** Summary of all the stimulus and behavioral variables used in this study for a representative individual fish. On the left: the entire duration of the experiment. On the right: close up of the trajectory and the relevant variables highlighted in the grey box. The four graphs show: (i) temperature experienced by an example fish, (ii) ΔTemperature (difference in temperature between current and last movement) experienced [color coding represents: worsening of conditions (in brown, away from physiological setpoint), improvement (in turquoise, toward physiological setpoint) or no change (in gray)], (iii) the absolute value of reorientation for each swim [color coding represents: turn (in red, when reorientation is higher than 30 degrees), or forward (in blue, when reorientation is lower than 30 degrees)] and (iv) the direction of turns, either left or right.

While the difference between the currently experienced and the animal’s preferred temperature is the fundamental drive for homeostatic navigation, this information alone does not provide a salient directional cue that could instruct directed behavior^9,10^. We reasoned that, similarly to chemotaxis, phototaxis and rheotaxis^11–14^, the simplest directional cue is given by the temporal changes in temperature due to animal self-relocation^3, 15^ (Figure 1 c). To improve its environmental condition, fish could thus rely on a simple ‘tumble and run’ strategy as used by bacteria^11, 16^. Thus, previously experienced temperatures could contextualize the current one and provide a basic notion of a worsening or improving context, driving an opposite behavioral output, like increased turning versus straight runs, independent of the absolute temperature^15^. A worsening context should increase the reorientation probability to change fish direction of travelling. On the other hand, an improving context, meaning getting closer to the homeostatic setpoint, should result in turning suppression in order to keep the current direction^3, 10, 11, 15, 17^ (Figure 1d). Since larval zebrafish swim in discrete bouts at an approximate rate of 1 Hz^18^ context evaluation should happen at the end of each swim event.

By dividing each movement in either a turn or a forward swim and by taking into account the temperature change experienced by the fish during the last motor action (see Methods), we were able to define three scenarios. An improving context (ICxt) when the temperature change was toward the fish’s setpoint independent of the absolute temperature experienced, a worsening context (WCxt) when the fish moved away from the setpoint and no context (NCxt) for an isothermal movement. Figure 1e shows an example fish moving in the arena together with all the relevant stimulus and behavioral variables we extracted from the trajectory. In the close-up of a behavioral sequence (Figure 1e right), we analyzed several periods where the animal experienced the same temperature but in two different contexts (orange arrow for WCxt and blue arrow for ICxt). In the WCxt (orange arrow) the reorientation probability is highly modulated by the sensory context (improving vs. worsening) such that turning frequency increases. Morever, as shonw in the bottom panel, fish tend to concatenate turns in the same direction where right turns are followed by additional right turns and vice versa.

By plotting the relative distribution of the absolute angle turned for all fish for WCxt and ICxt (Figure 2a) we confirmed our previous observation that temperature context influences reorientation. Worsening of experienced temperature led to higher turn angles with a higher probability of reorientation. For simplicity, we also generated a simple index that allowed us to quantify the amount of reorientation driven by sensory context in future analysis. In particular, we computed the difference between the turn fraction during WCxt and the turn fraction during ICxt. Again, we observed a significant difference (Figure 2b). Moreover, we noticed that the direction of turning was not stochastically decided during each movement^19^. In fact, when we quantified swim direction with respect to the previous turn event for the WCxt scenario, we found a high correlation, suggesting a form of memory used by the fish to persist in the previously chosen direction (Figure 2c)^19^. These data suggested that fish use a more sophisticated strategy than a simple ‘tumble and run’.

**Figure 2:**
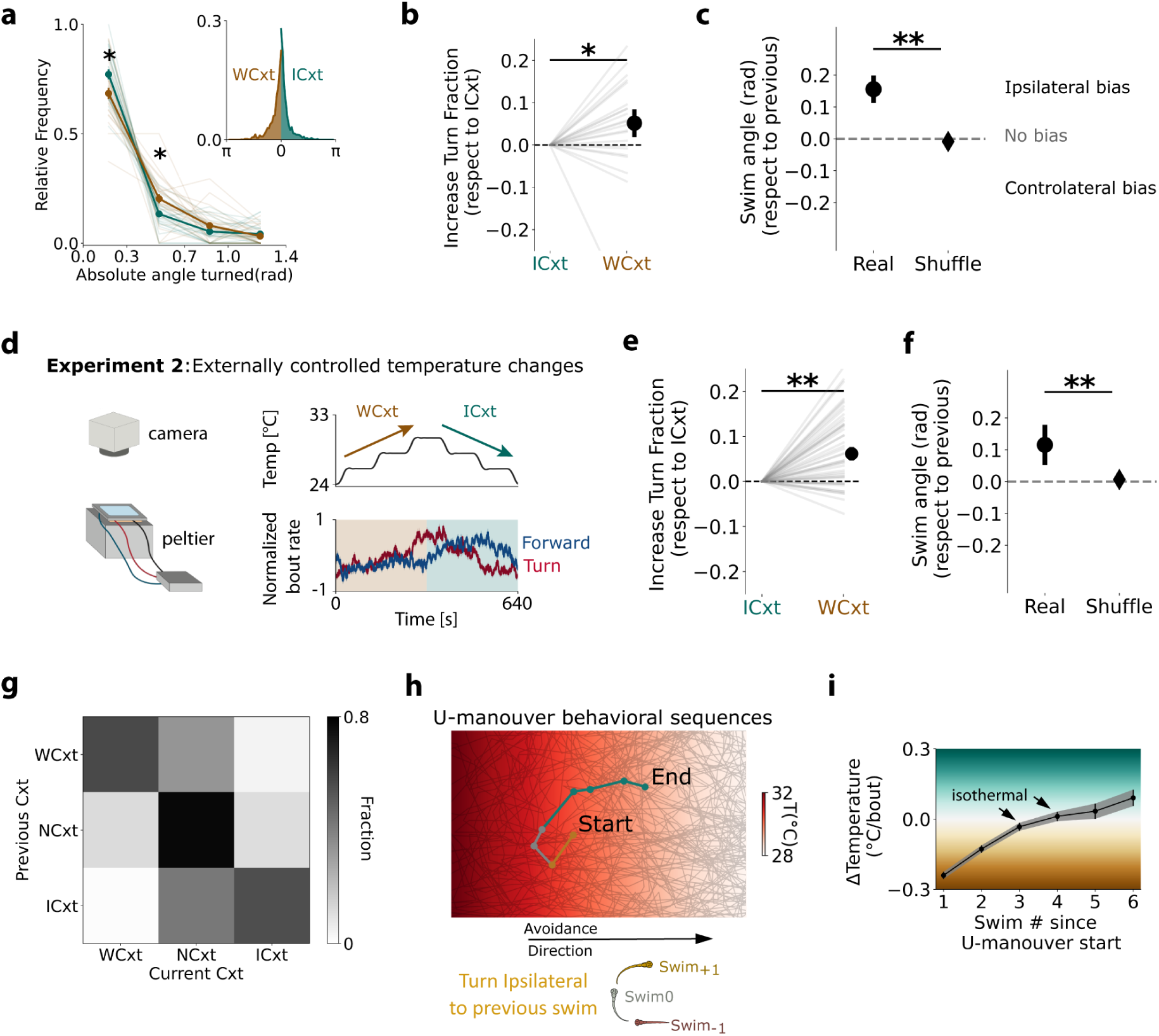
Homeostatic navigation is performed by combining an adirectional with a directional strategy. **a.** Relative distribution of the absolute angle turned when fish experienced a WCxt or an ICxt (mean ± standard error of the mean, Mann-Whitney non parametric test with Bonferroni correction for multiple comparisons). **b.** Normalized increase in turn fraction depending on sensory context (median ± standard error of the median, Mann-Whitney non parametric test). **c.** Direction of swim during WCxt according to the direction of the previous swim (mean ± standard error of the mean, Mann-Whitney non parametric test). Positive values imply that the swim tends to be in the same direction as the previous one. Diamond on the right is a shuffle obtained by assigning the sign for each value randomly. **d.** Left: Behavioral setup. Right Top: stimulus protocol for temporal gradient experiment. Right bottom: average normalized turning rate (red) and forward swim rate (blue) for n= 40 fish. **e**. Increase in turning fraction depending on sensory context for experiment 2, similar to **b** (median ± standard error of the median, Mann-Whitney non parametric test). **f**. Turn correlation upon WCxt for experiment 2, similar to **c** (mean ± standard error of the mean, Mann-Whitney non parametric test). **g**. Mean transition matrix for sensory context across fish population. **h**. Top: example U–maneuver performed during experiment 1. Bottom: sketch of a U-maneuver. **i**. Difference in temperature experienced by fish during the execution of U-maneuvers, starting from in a WCxt (mean ± standard error of the mean).

In order to extract the contribution of sensory history on the increase in reorientation probability and directional persistence, we challenged the fish with a step-like ramp ranging from 24 °C to 30 °C. Specifically, every two minutes, the temperature changed in the whole-arena by 2 °C or 0.016 °Cs^-1^ (Figure 2 d, see Methods). For these experiments, we developed a square-shaped, smaller arena, which allowed the temperature to change homogenously on the entire surface (Figure 2d left). The fish was, just as before, able to swim freely. Similar to previous experiments, analysis of the turn fraction (Figure 2d-e and Supplementary figure 2a-b, see Methods) showed that turns were indeed upregulated by sensory history (i.e. ICxt and WCxt) producing more reorientation maneuvers when the temperature was moving away from the preferred setpoint. Furthermore, analysis of the swim direction in the WCxt epoch confirmed also the high correlation in time of turning direction (Figure 2f)^19^. This result highlights the robustness and replicability of fish behavioral strategy across different setups and conditions.

In the large rectangular arena, we noticed, that due to the shape of the temperature gradient (Supplementary figure 1b, c), it is improbable for the fish to move from a WCxt to an ICxt in a single movement (Figure 2g and Supplementary figure 2c but see also Figure 1e). Consequently, such transitions usually take several swim events, where many of those movements do not provide any useful information about the change in temperature (NCxt). We hypothesized that fish could be able to use past changes in temperature and combine them with previous motor choices to persist in their behavioral program. We confirmed that fish were prone to reorient more and with ipsilateral swims when coming from a WCxt even if they did not experience a change in temperature during their last movement (NCxt) (Supplementary 2d and e). This phenomenon is invariant from the threshold used to define NCxt and therefore unlikely to arise from an analysis artifact (Supplementary figure 2f). In fact, the reorientation sequences produced U-shaped trajectories (U-maneuvers) commonly ending with the fish facing in the opposite direction, toward the preferred temperature (Figure 2h). In order to validate these observations, we extracted all the U-maneuvers started in a WCxt and computed the average temperature difference experienced by the animals during the execution of these behavioral sequences (see Methods). In line with previous quantifications, directional swims persisted in the direction defined during WCxt even in the absence of new sensory cues (Figure 2i, see swims 3 and 4 indicated by the black arrows).

To summarize our behavioral observations, we found that larval zebrafish perform homeostatic navigation by modulating the reorientation probability depending on the sensory history experienced while swimming. Going away from the homeostatic setpoint elicits movements leading to higher reorientation probability. Moreover, the direction of such reorientation maneuvers tends to be coherent, as in the observed U-maneuvers. We finally observed that these maneuvers persist even in the absence of immediate sensory directional cues (NCxt) suggesting the presence of a working memory.

### A whole-brain functional screen identifies brain regions modulated by sensory context

Our behavioral experiments indicate that zebrafish larvae regulate body temperature by making use of a sensory history and motor-dependent behavioral strategy. We took advantage of the optical transparency of larval zebrafish to perform a whole-brain functional screen using light-sheet microscopy (n=5, 6-8 dpf, *Tg(elavl3:H2B-GCaMP6s)*) to identify brain regions responsive to temperature, putatively involved in homeostatic navigation.

To control the temperature experienced by the fish while embedded and during imaging, we placed a small needle close to the fish head. This provided a constant flow of water through a multivalve system, which could also be heated or cooled down upstream of the tube by a Peltier element (see Methods, Figure 3a). Then, we first delivered the same stimulus protocol as in Figure 1d (Figure 3a top right) and in a small subset of fish (n=2) performed lightsheet whole-brain imaging. Behaviorally, we again observed a modulation of the reorientation probability depending on sensory context and direction for WCxt, replicating our freely swimming data (Figure 3a bottom right, Supplementary figure 3a, b, c and d). When we extracted neurons that reliably responded to the 3 trials (see Methods) we qualitatively observed multiple responding regions, such as the Olfactory Bulb (OB), Pallium (Pa), right Habenula (rHb), Preoptic area (PoA), Interpeduncular Nucleus (IPN) and Raphe nucleus (RphN) (Supplementary figure 3g).

**Figure 3:**
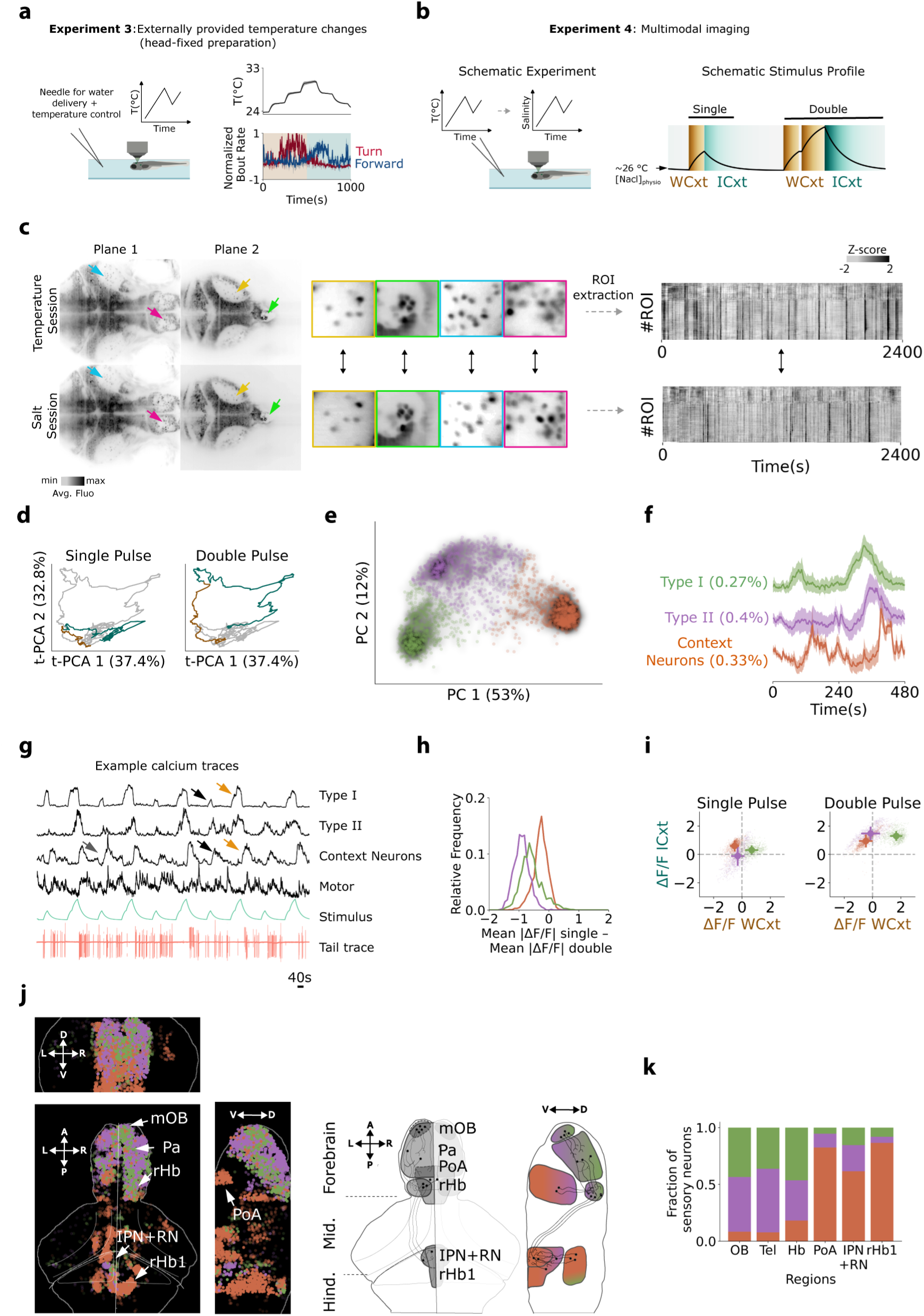
Identification of neurons encoding sensory context in a whole-brain screen. **a**. Left: Schematic representation of the head-restrained preparation for the lightsheet setup. Top right: protocol as in Figure 2d. Bottom right: average normalized turning rate (red) and forward swim rate (blue) for n=11 fish. **b**. Left: Schematic representation of the multimodal experiment. Right: Short protocol used for the lightsheet imaging experiments. **c**. From left to right: Anatomy of an example fish during the temperature and salt sessions. Yellow, green, cyan and purple arrows indicate four different patterns of neurons visible in both sessions. Raw activity of 10% of neurons (randomly selected) in a dataset for both temperature and salt session. Arrangement of the traces has been done according to the rostrocaudal position of selected ROIs. **d**. Temporal trajectories in PC space of the neuronal activity of reliable neurons, color-coded based on stimulus context (orange WCxt, blue ICxt). **e**. Projection onto PC space of all reliable neurons color-coded according to their cluster identity. **f**. Mean activity of each cluster (mean ± standard error of mean) and fraction of cells per group (tot n= 3270). **g**. Example raw traces from the three clusters and one motor ROI (all in black), the approximate stimulus profile (green) and the tail trace showing behavior (pink). **h**. Relative frequency distribution of the difference in mean fluorescence change during the single and double pulse stimulation for each cluster. Negative values imply that the fluorescence increased with increasing absolute temperature. **i**. Scatter plot of the average deviation from baseline fluorescence during single (on the left) and double pulse (on the right) split by sensory context (WCxt and ICxt) for the neurons in the three clusters (mean ± standard error of mean). **j**. Left: Anatomical distribution of reliable ROIs according to cluster identity. Difference panels depict different projections. mOB: medial Olfactory Bulb, Pa: Pallium, rHb: right Habenula, PoA: Preoptic Area, IPN+RN: Interpeduncular Nucleus and Raphe Nucleus, rHb1: Rhombomere 1. Right: Sketch of hypothesized synaptic connections based on activity maps and literature. **k**. Proportion of ROIs from the different clusters for different anatomical regions. OB: Olfactory Bulb, Tel: Telencephalon (Pallium+Subpallium), Hb: Habenula (Left + Right Habenula), PoA: Preoptic Area, IPN+RN (Interpeduncular Nucleus + Median Raphe + Dorsal Raphe), rHb1 (Rhombomere 1).

While this protocol elicited a strong and clear behavioral modulation, the relatively long duration of each trial (⁓20 minutes) was not optimal for an imaging experiment, where a large number of repetitions are beneficial. Moreover, observed neuronal responses could be linked to either temperature sensing or motor behavior. Given the strong correlation between the stimulus protocol (upward vs. downward temperature gradient) and the animal’s behavior (turn vs. forward), in this experiment they could not be disentangled. We therefore devised a shorter protocol with 5 repetitions of a shorter stimulus block. In each block, fish experienced an increase of +0.25 °C away from their preferred temperature (∼26 °C) before the temperature was brought back to baseline (single pulse). Then the increase was repeated, but this time the temperature was subsequently further increased to ∼ 26.5 °C (double pulse) (Figure 3b and Supplementary figure 3e). The rationale behind this protocol is to present the fish with a situation which can either improve after getting worse (single pulse, ICxt) or get worse again (double pulse, WCxt), thus mimicking a naturally occurring scenario that the fish encounters while swimming freely.

Temperature is not the only environmental factor, which poses a homeostatic threat and might lead to an asymmetric behavior when the animal is pushed away from its homeostatic setpoint^10^. Thus, we asked whether the fish brain encodes a representation of the stimulus history that generalizes across different sensory modalities, regardless of the stimulus identity. Therefore we recorded the brain activity during transient increases in salinity (salt session), which can be equally threatening to the animal’s physiology^10, 20^. Importantly, we were able to match the stimulus profile of the salt session with the temperature session (Supplementary figure 3e and Methods), to properly compare brain responses independent of sensory modality. In addition, we carefully matched single neuron identity across the two sessions (Figure 3c, see Methods) and extracted their fluorescence throughout (randomly extracted 20 % ROIs Figure 3e right). For each fish we extracted 44,772 ± 7,090 (mean ± std.) ROIs spanning the forebrain, midbrain, and parts of the hindbrain until rhombomere II.

We first focused our analysis on the temperature session. In particular, we employed an unbiased approach to screen for neurons responding to different properties of the stimulus. For each neuron we computed a reliability index across trials and across individuals (see Supplementary figure 3g and see Methods), we selected only “reliable neurons” using a fixed standard threshold criterion for all fish and computed the trial-triggered average (TTA) for all included neurons. We then performed principal component analysis (PCA) on the time dimension. When we projected the neuronal activity to the first two temporal principal components (cumulative variance explained 70.2 %, Figure 3d), neurons showed different responses in the single pulse compared to the double pulse. Whereas as the first principal component, t-PC1, appeared to separate activity based on the context the second principal component t-PC2 conveyed information about the absolute temperature. To better understand what type of neuronal responses were responsible for the context-dependent activity, we performed PCA in the other dimension so that each dot in the 2D space shown in Figure 3e represented a particular neuronal response (cumulative variance explained 65% variability, Supplementary figure 4a). Since 3 clusters were clearly observable, we performed k-means clustering (k=3, Figure 3e). The number of clusters was further checked using the Davies-Bouldin score (see Methods and Supplementary figure 4b).

The three clusters corresponded to three populations of neurons: a population following the stimulus profile (type I), a population showing an increase in activity upon temperature rises with a longer rise constant (type II) and a population that showed a reduction in baseline fluorescence when temperature increased (WCxt) and an increase in fluorescence when temperature decreased (ICxt) (type III) (Figure 3f). As shown by the raw traces in Figure 3g, the responses in the last group do not arise from an averaging artifact or motor activity. They are already visible within individual fish at the single trial level and are dissociated from purely motor-related activity. Neurons of type III are, therefore, well-suited to inform the fish on whether the temperature context is worsening or improving (context neurons). Moreover, when we computed the difference in peak fluorescence between the single and the double pulse within the stimulus block, we found that activity in type I and type II neurons were modulated by absolute temperature. Conversely, activity in context neurons showed minor modulation, suggesting that responses in this cluster did not scale with stimulus intensity (Figure 3h, see also raw traces in Figure 3g, black and orange arrows). The same observations were recapitulated when we computed the average deviation from baseline, split by context (WCxt or ICxt) in the single or double pulse periods: the type III neurons showed very similar activity in both pulse conditions (Figure 3i).

Importantly, when we looked at the anatomical distribution of the three clusters we noticed that Type I and II neurons were enriched in rostral regions like the Olfactory Bulb (OB), Pallium (Pa) and right Habenula (rHb) while context neurons were found in other regions like the Preoptic Area (PoA), anterior Hindbrain (aHb), Interpeduncular Nucleus (IPN), and Superior Raphe (RN). Most of the aforementioned regions are known to be strongly connected (Figure 3j)^21–24^. In fact, we observed a functional gradient along the OB-Pa-Hb-IPN-RN pathway, with upstream regions mostly responding to absolute temperature and downstream regions, together with the PoA, mostly tracking the stimulus context (Figure 3k).

Having identified several brain regions enriched in context neurons, we next wanted to find putative pre-motor areas that could use this information to influence behavior. Therefore, we searched for cells that were reliably activated by either forward, left or right swims (Figure 4a and Methods). In line with other reports^25–29^, we observed neural correlates of motor activity around the nucleus of the medial longitudinal fasciculus (nMLF) and hindbrain regions. We also examined the localization of motor-related signals in brain regions that were previously identified as crucial for tracking stimulus features. We noticed that regions with a higher number of context neurons tended to also have more motor correlates (Figure 4b). This activity, locked to swimming events, could be directly instructive or could represent a short-term memory trace of what the animal did in the immediate past. In the latter case, this representation could provide a motor context required for the U-shaped behavioral sequences that we observed in our freely moving experiments, where the turning direction of subsequent bouts was highly correlated (see Figure 2i). We reasoned that a potential neural substrate for such motor memory should lag with respect to a canonical premotor neuron and display more persistent activity. In order to find any temporal progression of neural activity following a turn, we clustered turn-responding neurons depending on the timing of their peak activity (Supplementary figure 4c) and then computed the average activity in each cluster (Supplementary figure 4d, see Methods). As shown by the anatomical maps of the different timing clusters (Supplementary figure 4e), turn-related activity elicited a temporal progression of different neuronal populations. Fast neurons are localized contralateral with respect to turn direction and in a diffuse area of rhombomere II while slow neurons are ipsilateral and localized in a small medial region of the aHb^25, 30, 31^.

**Figure 4:**
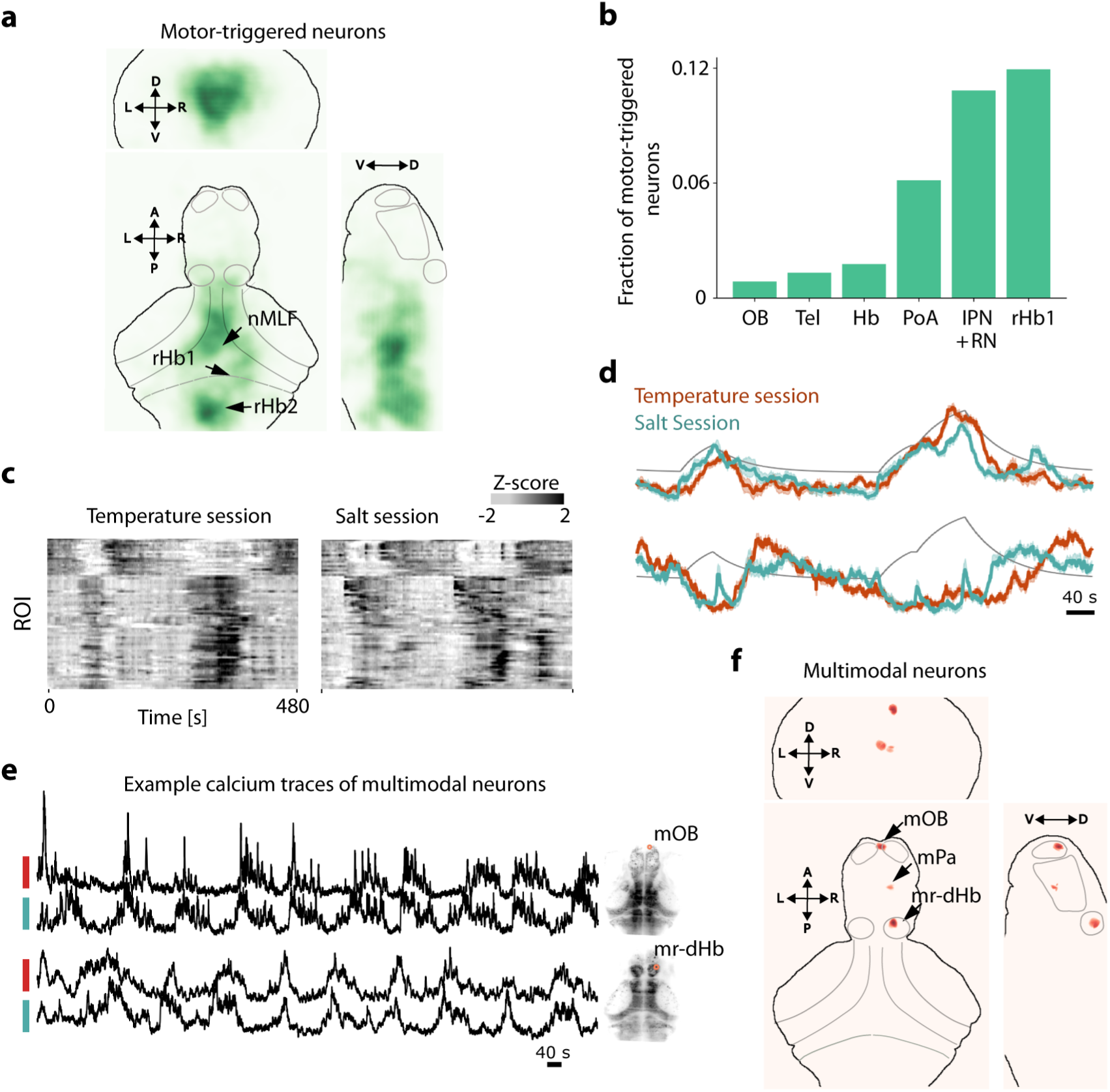
A small population of dorsal Habenula, medial Olfactory Bulb and Pallial neurons respond similarly to temperature and salinity changes. **a**. Anatomical density distribution of motor-triggered ROIs. Selected cells can be either forward, left or right-swim tuned. **b**. Fraction of motor-triggered ROIs for each sensory-identified region. **c**. Trial trigger average of all multimodal neurons found during temperature (left) and salt (right) sessions. **d**. Mean of the multimodal neuron clusters (k=2, mean ± standard error of the mean). **e**. Raw activity of two example multimodal neurons (one from the olfactory bulb and one from the habenula) in black for both the temperature and the salt session. **f**. Anatomical density distribution of multimodal neurons for the three projections mOB: medial Olfactory Bulb, mPa: medial Pallium, mr-dHb: medial nucleus of the right dorsal Habenula.

Finally, we looked for multimodal neurons with similar stimulus response profiles for both the salt and the temperature sessions. We cross-correlated the TTAs of reliable neurons in both sessions and then computed a p-value based on circular shift shuffle versions of such traces (see Methods). We selected neurons with significant p-value<0.025 (Figure 4c and Supplementary figure 4f) and clustered them into two groups (Figure 4d). This analysis revealed neurons with similar dynamics upon temperature and salinity changes (Figure 4e). While all the previously identified sensory regions had neurons responding during both sessions (Supplementary figure 4g), only the medial nucleus of the right dHb (mr-Hb), the mOB and a small region in the Pa, had multimodal neurons (Figure 4f).

In summary, our whole brain analysis confirmed that temperature is sensed by neurons located in the olfactory system^32^, probably among other regions that were not imaged in this study such as the trigeminal ganglion^33^. This information is then broadcast to hypothalamic regions such as the PoA and from there to the dHb-IPN-RN system. The PoA was particularly enriched in temperature specific context cells^34^ suggesting a role in thermoregulation, just as in mammals^32, 35–39^. Regarding the dHb-IPN-RN system, the situation is more complex. Although responses in the dHb are mainly sensory, responses in the IPN, the main target of the dHb, were enriched in context and motor-modulated cells. The Hb is also known to receive direct input from the Pa and the PoA. Finally, the Hb was also particularly enriched with neurons that generalized across modalities. Overall, suggesting that both the PoA and the Hb could be involved in conveying the sensory context during homeostatic navigation.

### PoA and dorsal Habenula ablations impair different aspects of homeostatic navigation

In our whole brain screening approach, we identified the PoA and dHb-IPN-RN as candidate pathways to provide information about the stimulus context. Considering the role of the PoA in thermoregulation and the fact that it sends projections to the Hb, we hypothesized that they could both be involved in homeostatic navigation. To test this, we ablated the PoA or the dHb and analyzed the behavior of the fish in the spatial gradient arena.

Bilateral ablations of the PoA, as functionally identified in our previous light-sheet experiments, were performed with a Ti-Sapphire laser at 5 dpf (Figure 5a, Supplementary figure 5a). As a control, we targeted part of the medial Optic Tectum (OT), since this region was poorly activated by temperature stimuli (n=20 for both groups, see Methods). After the ablation procedure, fish were left to recover for 40-48 hours and tested at 7 dpf. Ablation of the PoA produced a higher coefficient of dispersion in the thermal gradient at the end of the experiment compared to controls, suggesting an impairment in staying near their homeostatic setpoint (Supplementary figure 5b and c, see Methods). To test this hypothesis, we computed the same increase in turn fraction used for wild-type fish. Upon PoA ablation, fish showed a significant impairment in the modulation of reorientation probability depending on sensory history (ICxt vs. WCxt) with respect to controls (Figure 5b). The absence of such modulation is also reflected in the reduction of turn fraction, compared to controls, during the entire experiment (Supplementary figure 5d). Notably, also the directional persistence to change orientation through U-maneuvers when the fish experienced a WCxt was impaired (Figure 5c). These results suggest an essential involvement of the PoA in driving reorientation behavior to navigate toward the homeostatic setpoint.

**Figure 5:**
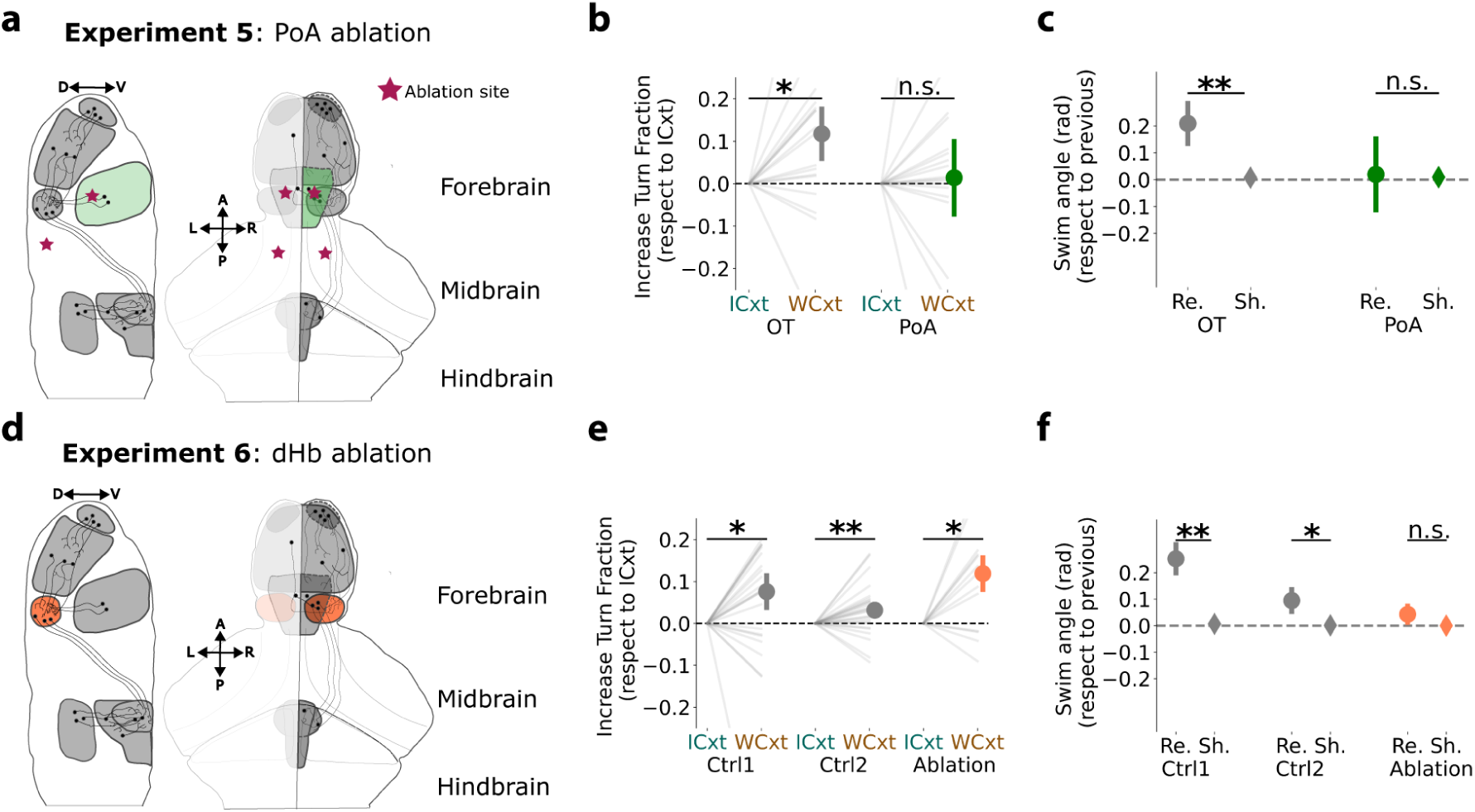
PoA and dHb ablations impair different aspects of homeostatic navigation. **a**. Sketch of 2-photon ablations targeting the PoA (depicted in green). Stars are the chosen ablation sites. The control ablation is in the optic tectum (OT) (see text). **b**. Increase in turn fraction depending on sensory context. Left: control OT ablation, right: PoA ablation. (median ± standard error of the median, Mann-Whitney non parametric test). **c.** Turn correlation upon WCxt for the control and ablation groups (mean ± standard error of the mean, Mann-Whitney non parametric test). **d**. Sketch of chemogenetic ablations targeting the dHb (depicted in orange). **e**. Increase in turn fraction depending on sensory context. Left: genetic control, middle: treatment control, right: dHb ablation (median ± standard error of the median, Mann-Whitney non parametric test). **f.** Turn correlation upon WCxt for the two controls and the ablation group (mean ± standard error of the mean, Mann-Whitney non parametric test).

We next used the transgenic *Tg*(*Gal4:16715; UAS:Ntr-mCherry*) line to ablate the dHb at 5 dpf using a chemogenetic approach (see Methods). This line restricts the expression of Nitroreductase (Ntr) to the dHb (Figure 5d and Supplementary figure 6a, b). As control groups, we used a genetic control and a treatment control (n=25, 23 and 14 for genetic control, treatment control and ablation group respectively, see). To our surprise and in contrast to the PoA ablations, even though the group dispersion coefficient was higher for dHb ablated fish compared to controls (Supplementary figure 6d), dHb ablated fish were still able to modulate their reorientation probability based on sensory history (Figure 5e). However, when we tested the use of a working memory for sensory history before the last movement, namely when the fish had access to intermittent sensory cues (e.g. WCxt→NCxt or ICxt→NCxt), the increase in reorientation probability was lost in the ablated group (Supplementary figure 6f). Instead, we observed a general upregulation of reorientation maneuvers in the ablated group (Supplementary figure 6e). Finally, similarly to the PoA ablations, the correlation in swim direction (U-maneuvers) was impaired in Hb ablated fish compared to controls (Figure 5f).

These results confirm our findings using calcium imaging and reveal that the PoA and the dHb are both involved in homeostatic navigation, and they jointly support it. On the one hand, the PoA compares the distance from the setpoint before and after a swimming event and triggers reorientation when the fish is in a WCxt. It also conveys this information to the dHb. On the other hand, the dHb uses the inputs from the PoA to produce coherent steering trajectories enabled by a working memory of the sensory history needed especially in shallow temperature gradients where individual movements do not change the experienced environmental temperature. Thus, the dHb supports the fish in directional navigation in the absence of constant new information after each movement.

## Discussion

In this study, we have demonstrated that larval zebrafish, an ectotherm, can navigate a shallow thermal gradient to locate its preferred environmental temperature and thereby achieve temperature homeostasis. This contrasts with endotherms such as mammals who rely on autonomic responses in addition to behavioral adjustments^6^. Importantly, our data reveals that temperature homeostasis in an ectotherm still relies on the PoA, the internal thermostat of the mammalian brain. Moreover, our results suggest that the dHb, connected to and downstream of the PoA^24^, enables precise temperature homeostasis in environments with slow or minor temperature changes through working memory.

More specifically, the strategy larval zebrafish use involves comparing the temperature before and after a swimming event and by biasing their subsequent behavior based on whether the new temperature approached their preferred setpoint (ICxt) or was further away from it (WCxt). In the first case, the fish responds with more forward movements, while in the latter case, they reorient, in order to change their direction of travel. This non-directional strategy is widely used across the animal kingdom when navigating many different sensory gradients^3, 10–14, 16, 17, 40^ and has been referred to as behavioral hysteresis^15^.

In our experiments, we also observed that fish tend to concatenate reorientation events, where a turn is likely followed by another one in the same direction^41^. Finally, we found that when a swim led to neither an improvement (ICxt) nor worsening (NCxt), fish were still able to leverage their memory of the sensory history before the last movement and combine it with the knowledge of the previous movement direction to generate coherent U-maneuvers (Figure 6, top).

**Figure 6:**
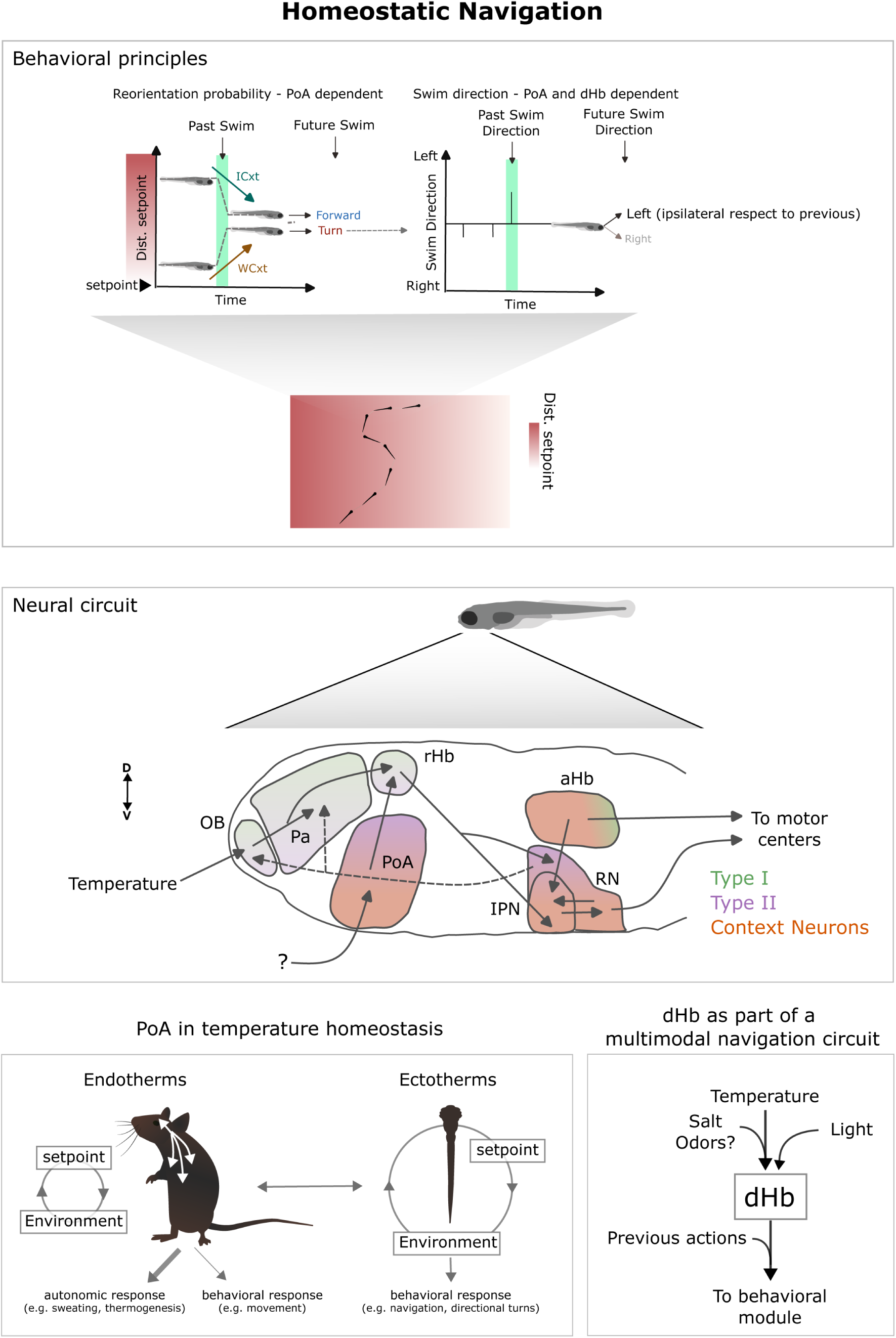
Homeostatic navigation. Top: Behavioral principles underlying homeostatic navigation in larval zebrafish uncovered by this study. Fish use two complementary strategies: (i) an increase in turning (reorientation probability) when experiencing a WCxt (being pushed away from the homeostatic setpoint) coupled with (ii) turning in a persistent direction during this WCxt. The first strategy is PoA dependent whereas the second depends on both the PoA and the dHb. Middle: Neural circuitry involved in homeostatic navigation. Neurons representing the three main different stimulus features are represented in different colors and the anatomical shading shows the fraction of response types in each region. Bottom left: Comparison of the role of the PoA in endotherms vs ectotherms. Bottom right: the dHb as a structure through which different sensory modalities are funneled to generate a common homeostatic navigation strategy.

Moreover, the fact that similar sensorimotor strategies have also been described in different types of temporal and spatial taxis assays^10, 14, 42, 43^ raises the hypothesis that fish can also have an abstract representation of context, independent of the sensory modality. Fish could then use a unique navigational system informed by different sensory modalities rather than independent parallel unimodal circuits.

Overall, our behavioral data suggest that homeostatic navigation is more complex than a simple reflexive stimulus-response behavior. The mechanisms involved in this process go beyond the moment-by-moment estimate of the sensory history with respect to the homeostatic setpoint. We propose that even when new sensory information is absent, fish can still use their past motor choices to guide an internally generated motor program. This program has a directional component and includes several swimming events. Thereby, fish can achieve precise temperature regulation also in slow or minimally changing temperature environments.

### The neural circuit enabling homeostatic navigation

Taking advantage of whole-brain functional imaging, we identified brain regions responding to temperature changes in an open-loop virtual gradient assay (Figure 6, middle). Temperature responses predominantly fell into three functional types. Type I and type II neurons responded to the stimulus profile but with different dynamics while type III tracked stimulus change (context neurons). The activity of context neurons was bidirectionally modulated, with a reduction from baseline during WCxt and increased activity during ICxt. Their activity was not influenced by stimulus intensity. We also observed a functional gradient along the rostro-caudal anatomical axis. Neurons of Type I and II were mostly localized in the forebrain of the fish. The medial Olfactory Bulb (mOB) and the Pallium (Pa) had almost exclusively Type I and Type II cells while the right Habenula (rHb) showed a mixture of Type I, II and context neurons. A possible interpretation as to why we see activity in regions traditionally linked to olfaction^33^ could be that, in a water medium, temperature and chemicals (such as odors) tend to have similar spatiotemporal statistics and is therefore advantageous to exploit the same sensory circuit for both. Interestingly, temperature sensing in olfactory circuits has also been found in other animals^44–46^. This interpretation is supported by reports of recruitment of larval zebrafish olfactory regions upon exposure to salt^10^, changes in pH^20^ and carbon dioxide^47^.

On the other hand, context neurons were predominantly – with the notable exception of the Preoptic Area (PoA, see below) – found in the fish midbrain and hindbrain in regions like the anterior hindbrain (aHb), the Interpeduncular Nucleus (IPN) and Raphe Nuclei (RN). Interestingly, bidirectional modulation of neuronal activity in rhombomere I has also been recently reported in an OMR-based decision-making paradigm involving leftward and rightward visual motion stimuli^29^. This commonality raises the possibility that neurons in these regions are not involved in sensory processing per se but in the selection of two opposite behavioral strategies based on sensory context.

Most of the regions highlighted are also known to be strongly connected. Specifically, neurons in the mOB project to both the Pa and the rHb, often via collaterals of the same neuron^23^. Neurons in the medial portion of the Pa project to the rHb^21, 22^, together with the PoA^24^. The Hb is, then, well known to send glutamatergic input to the IPN through the Fasciculus Retroflexus^21, 22^. In particular, neurons in the rHb selectively project to the ventral portion of the IPN with axons wrapping around this structure. Finally, the IPN is a GABAergic nucleus projecting to RN and receiving inputs from the aHB^21, 48^. Strikingly, the RN sends axons to more caudal rhombomeres as well as feedback projections back to forebrain regions such as the OB and Pa^49^. We, therefore, suggest that these regions are part of a brain-wide network supporting homeostatic navigation.

### Preoptic Area – a homeostat in ectotherms?

We investigated a potential role of the PoA in larval zebrafish thermoregulation as this structure is critical in mammals for maintaining a stable body temperature and it was particularly enriched in context neurons. Following the bilateral ablation of this region, we found a deficit in the test group in reaching thermal homeostasis. Fish failed to modulate their reorientation probability depending on the sensory context with respect to controls.

Unlike the dHb pathway (mOB-rHb-IPN-RN), less is known about PoA anatomy in larval zebrafish. Since the main targets of the OB are the rHb, Pa, Posterior Tuberculum and the Ventral Nucleus of the Ventral Telencephalon^23^ it is unclear whether the PoA receives temperature information from the OB, Pa, or other structures such as the terminal nerve^22^, trigeminal or dorsal root ganglion^33^.

These results are consistent with the previously proposed role of the PoA in larval zebrafish in detecting temperature changes^33^ and reacting to various types of physiological stressors^34, 50, 51^. However, our results expand those findings by suggesting a role of the PoA in thermoregulation by evaluating moment-by-moment the sensory history with respect to the homeostatic setpoint and conveying to the dHb information to persist in the chosen motor program and achieve temperature homeostasis.

We, therefore, propose that the PoA plays an evolutionary conserved function in regulating body temperature across vertebrates (Figure 6, bottom left). We suggest that this function has adapted to the physiology and the nervous systems of different organisms. In mammals, which have both autonomic and volitional strategies to cope with thermal stress, the PoA integrates peripheral and central temperature information^6, 52^ to increase thermogenesis and physical activity^39^. Conversely, here we provide evidence that the PoA in larval zebrafish directly triggers active navigation as an efficient homeostatic control mechanism in an animal without internal homeostatic mechanisms.

Our findings fit within a broader framework suggesting that the PoA and the hypothalamus, in general, can influence behavior at faster timescales than previously thought^53–56^. In the future, it would be interesting to understand if particular cell subtypes in the PoA are specifically involved in thermal navigation like has been shown for nocifensive behavior^50^ and the interplay between the PoA and the Hb-IPN-RN pathway.

### Dorsal Habenula – working memory for precise homeostasis

Except for the PoA, all other brain regions in our whole-brain screen which were enriched in context neurons, namely the IPN and RN, are located in the midbrain-hindbrain boundary, are mono-synaptically connected to each other and receive inputs from the Habenula (Hb).

In our imaging experiments, the right dorsal Hb (rdHb), together with a small region in the medial Olfactory Bulb (mOB) and in the medial Pallium (mPa), responded reliably to both increases in temperature and salinity with similar response profiles. These results are in line with other reports highlighting the contribution of the dHb to other types of sensory-driven navigation behaviors^10, 43, 57–59^. We suggest that the dHb could provide a more general estimate of environmental context to mediate an

appropriate behavioral response. In this scenario, different sensory modalities would converge onto a single navigation circuit to instruct a shared behavioral strategy. The dHb was also the first station in the above-described pathway to have context neurons and to show no modulation due to motor activity.

Contrary to our expectations, dHb ablation did not impair modulation of reorientation probability based on sensory context (i.e. ICxt and WCxt). However, the coherence in turn direction during WCxt was abolished. Moreover, we found an impairment in short-term working memory. These findings suggest that the dHb integrates contextual information on a timescale longer than a single movement and is involved in the directional component of homeostatic navigation (i.e. left vs. right).

In fish, the dHb and vHb (ventral Habenula) are the homologs of the medial and lateral habenula (MHb, LHb) in mammals^60^. The dHb conveys information coming from the limbic forebrain to the interpeduncular nucleus (IPN), while the vHb is directly connected to the median raphe (MRN). In adult and juvenile zebrafish the ability to switch behavioral strategies in different types of learning tasks is affected by manipulations of the dHb-IPN pathway^61, 62^ while in larvae the same pathway mediates odor attraction^57, 59^, CO_2_ and salt avoidance^10, 58^ and phototaxis^43^. The Hb has also been previously linked to temperature sensing in larvae^33^. It has also been proposed that the dHb is a multimodal trigger network associated to state transitions during foraging behavior^63^.

Previous studies interpreted the behavioral impairment following dHb ablation in taxis assays as a shift in stimulus preference^43, 57, 59^ and concluded that this structure conveys information about the stimulus valence to a behavioral module. The seeming discrepancy between these conclusions and ours can be reconciled by the difference in the stimulus landscape used and the amount of contextual information available to the animal in the arena. Here, we complement this model and propose that, rather than absolute valence, the dHb conveys a contextual relative valence signal computed from the animal’s recent past which, then, modulates directional choices (Figure 6, bottom right). This mechanism is particularly useful if the sensory landscape presents areas with few contextual cues like isothermal regions in shallow or slow changing gradients or split field arenas^43^. dHb’s role in homeostatic navigation might be a functional precursor of more complex cognitive abilities observed for this region in zebrafish adult stage and mammals such as valence^64^ and direction-based decision-making^61^ and strategy switching^60, 62^.

In future works, it would be interesting to understand if the dHb pathway is actively involved in sensorimotor transformation and/or acts as a global modulator, mostly through the serotoninergic and dopaminergic system, of brain dynamics^48, 63^. Lastly, the activity observed in the aHb might be an important part for the generation of persistent re-orientational sequences^48^.

In summary, our study shows that in a vertebrate model thermal regulation is achieved by the synergy of two brain regions, the PoA and dHb, which jointly through different contributions support homeostatic navigation.

## Acknowledgements

RP and IGK were funded by the Deutsche Forschungsgemeinschaft (DFG, German Research Foundation) as part of the SPP 2205 – project 430156228. In addition, RP was supported by grants to from the Volkswagen Stiftung Life? Initiative and by the German Research Foundation (DFG) under Germany’s Excellence Strategy within the framework of the Munich Cluster for Systems Neurology (EXC 2145 SyNergy, identifier 390857198). The authors would like to thank Thomas Frank, Florian Richter and Luigi Petrucco for the insightful discussions and comments on the manuscript and the Grunwald Kadow and Portugues labs for their input and encouragement.

## Methods

### Zebrafish husbandry

All procedures related to animal handling were conducted following protocols approved by the Technische Universität München and the Regierung von Oberbayern. Adult zebrafish (Danio rerio) from Tüpfel long fin (TL) strain were kept at 27.5-28 °C on a 14/10 light cycle, and hosted in a fish facility that provided full recirculation of water with carbon-, bio- and UV filtering and a daily exchange of 12% of water. Water pH was kept at 7.0-7.5 with a 20 g/liter buffer and conductivity maintained at 750-800 μS using 100g/liter. Fish were hosted in 3.5 liter tanks in groups of 10 to 17 animals and fed the adults with Gemma micron 300 (Skretting USA) and live food (Artemia salina) twice per day and fed the larvae with Sera micron Nature (Sera) and ST-1 (Aquaschwarz) three times a day.

All experiments were conducted on 5-7 dpf larvae of yet undetermined sex. The week before the experiment, one male and one female or three male and three female animals were left breeding overnight in a Sloping Breeding Tank or breeding tank (Tecniplast). The day after, eggs were collected in the morning, rinsed with water from the facility water system, and then kept in groups of 20-40 in 90 cm Petri dishes filled with Danieau solution 0.3x (17.4 mM NaCl, 0.21 mM KCl, 0.12 mM MgSO4, 0.18 mM Ca(NO3)2, 1.5 mM HEPES, reagents from Sigma-Aldrich) until hatching and in water from the fish facility afterwards. Larvae were kept in an incubator at 28,5°C and a 14/10 hour light/dark cycle, and their solution was changed daily. At 4 or 5 dpf, animals were lightly anesthetized with Tricaine 440 mesylate (Sigma-Aldrich) and screened for fluorescence under an epifluorescent microscope. Animals positive for GCaMP6s and mCherry fluorescence were selected for the imaging experiments.

### Transgenic animals

The Tuepfel long-fin (TL) wild-type strain was used for freely swimming behavioral experiments. The nacre *(mitfa−/−*, lacking melanophores) transgenic zebrafish lines Tg(elavl3:GCaMP6s+/+), labelling all the neurons, and *Tg(16715:GAL4VP16);Tg(UAS:NTR-mCherry)),* labelling the dorsal part of the Habenula, were used respectively for functional imaging experiments and chemogenetic ablations.

### Experimental Setups

#### Freely swimming rectangular arena (large arena)

The custom-made arena consists in a rectangular pool (200 × 40 × 3.5 mm^3^) (Supplementary figure 1a) made of aluminum for homogenous heat dispersal. The surface of the plate is taped, before each batch of experiments, with white tape (Tesa) to increase contrast and allow online tracking of fish. At both ends of the arena two Peltier modules (digikey, TEC-40-39-127) were fixed with thermal tape or thermal glue (Conrad Electronic). The arena is placed on a metal block which dissipates heat from the Peltier elements and whose temperature is constantly monitored (RS Components, Nr. 706-2743). The temperature is constantly monitored at both ends of the pool with waterproof thermocouples type Pt100 (RS Components, Nr.762-1134). Temperature was controlled by two independent TEC controllers (Meerstetter, TEC-1091). After about 5 min, the temperature of the water at both sides of the pool reaches the target temperatures (25 °C and 33 °C), which then remains constant over time. Water level is usually between 3-4 mm. Freely swimming larvae are monitored using a Ximea camera (MQ022MG-CM) at 110 fps, coupled with a macrolens (Navitar). The whole apparatus is placed in a light-tight box, illuminated with a homogeneous IR light emitted by a LED panel (Wilktop) placed on both sides, above the arena, spanning the entire small side of the setup.

#### Freely swimming square arena used for long temporal gradient experiment (small arena)

The arena consists of a square pool (40 × 40mm) (Figure 2d left) made of aluminum filled to a depth of 3-4 mm. A single 36 W Peltier element (50 x 50 x 3.5 mm) (digikey, TEC-40-39-127) is sandwiched between the heat sink and the arena and fixed with thermal tape (Conrad Electronic). Temperature is, therefore, changed homogenously in the entire volume of water (water level 3-4 mm). The Peltier is driven by a single TEC controller (Meerstetter, TEC-1091) and temperature is constantly monitored with a Pt100 thermocouple placed in the center of the arena (RS components, Nr. 891-9145). Homogenous illumination is provided from the side using IR light emitted by LED stripes (Solarox). Fish are tracked at 150 fps with a Ximea camera (MQ013MG-ON), coupled with a lens (Edmund Optics).

#### Perfusion system for head-embedded preparation under lightsheet microscope

We implemented a system where the head of the tethered fish is targeted with a constant flow at a rate of 1.5 ml min−1 of filtered fish water delivered through a needle (19 gauge) placed in front of the fish at an angle of 20-30° via a gravity-based 4-channels perfusion system (Figure 3a left). The needle is connected to an in-line solution heater/cooler (Warner Instruments, SC-20) to precisely regulate the temperature. Excess heat produced by the SC-20 Peltier is dissipated through a liquid cooling system (Koolance, Cat. EXT-1055). Water is constantly removed from the chamber to avoid overflowing using a small peristaltic pump controlled with an Arduino. The procedure ensures no changes in the water level (important for functional imaging purposes) and doesn’t affect the behavior of the fish.

The desired temperature is set through a single channel temperature controller (Warner Instruments, Cl−100) externally triggered by the computer handling the behavior protocol through a LabJack series U3-LV (LabJack). The controller also allows reading online the temperature inside the Heater/Cooler element and inside the chamber with a thermocouple which was placed immediately in front of the needle.

For salt experiments, we kept the temperature of the water flowing in the chamber at room temperature (24 °C) and we open/close a second valve connected to a reservoir with fish water at higher salt concentration (40 mM NaCl added to fish water). The switch is mediated by solenoid valves pinching the tubing and whose state is controlled by a ValveLink 8.2 perfusion controller (Automate Scientific) timed with the behavior protocol through an Arduino 1 (Arduino). A perfusion pencil tip combining 4 tubes into a single tip (AutoMate Scientific, 04-08-250) placed between the solenoid valves and the inline heater/cooler (switched off for this particular experiment) ensures rapid liquid volume exchange.

#### Behavioral Experiments

All the experiments where behavior was recorded were run from 10:00 h to 20:00 h. Since all experiments were carried out in darkness, we removed the fish from the incubator at least 2 hours before testing.

#### Spatial gradient in the large arena

For controlling the thermal gradient stability, on top of having the temperature constantly monitored at both ends of the pool by waterproof thermocouples, we divided the chamber into 10 bins of 2 cm each and we measured the temperature in each bin with a digital thermometer at the beginning and at the end of each experiment. Individual fish were then transferred to the arena and their behavior was monitored in the gradient for 15 minutes. Larvae were placed in the pool after the target temperatures (24 °C and 33 °C) were reached. The warmer side was randomized across experiments.

#### Long temporal gradient in the small arena and for head-restrained preparation

This protocol was presented to both head-restrained larvae under the lightsheet microscope and freely swimming larvae in the small square pool. The experiments lasted in total 60 minutes. The temperature was changed in the whole arena from 24 °C to 30 °C and then back to 24 °C with 2 °C steps lasting 2 minutes each. Each fish was presented with three repetitions of this ramp stimulus. At the end and at the beginning of each ramp the fish had 5 minutes where the temperature was kept constant at 24 °C. The temperature was monitored online.

#### Multimodal short temporal gradient for head-restrained preparation

Each head-restrained larval zebrafish we tested with this protocol was presented with a salt followed by a temperature session or vice versa. Before starting the experiment, fish were placed under the microscope with the laser on and fish water flowing for five minutes to let them habituate. The protocol comprised 5 repetitions of the same stimulus block. Each stimulus block lasted 480 s. During the experiment, room temperature was kept at 24°C, and the water was flowing at a low rate (<1.5 mL/minute). The overall protocol is described below. After 40 seconds, there induced a change in temperature or salinity (depending on the session type) for the next 40 seconds. For the temperature session, the peltier element was set to a target temperature of 27°C, resulting in an effective temperature increase of +0.2 °C while for the salt session we opened a second valve and released a high salinity solution. Afterward, there was a 90 second pause to allow the temperature or salinity to return to the baseline level. The second part of the experiment began with 50 seconds of temperature increase (with target temperature of 27 °C) or 40 seconds of an increase in salinity followed by a 10-second pause. Then, for the next 40 seconds, the target temperature of the peltier was set to 29 °C (effective temperature increase of +0.4 °C), or there was another 40-second increase in salinity. The block ended with 140 s of pause (experiment 4, Figure 3b right). The length of the pulses and pauses was carefully chosen to obtain the same stimulus profile across modalities and to not have an accumulation of temperature and salinity in the chamber over time (Supplementary figure 3e, f). To this end, we previously measured the conductance in water with an Arduino and we matched change in medium conductance (proportional to salinity concentration) from baseline to what we recorded with the thermocouple in the chamber. Randomization of double and single pulses was not possible due to accumulation phenomena in the case of two double pulses coming one after the other.

### Lightsheet functional imaging

#### Lightsheet microscope

For our experiments we used a custom-built microscope with two excitation scanning arms placed at 90° from each other. A laser beam coming from a 473 nm laser source (Cobolt) is split and directed into the two arms. In both arms, the laser beam gets expanded by a telescope before being focused through a glass coverslip on the fish. By scanning at 800 Hz the beam on the horizontal plane we generated the excitation lightsheet. The brain of the fish is targeted from the side and from the front giving access to the entire brain at a single-cell resolution. The emitted fluorescence was collected through a water immersion objective (Olympus, Japan), mounted on a piezo (Piezosystem Jena, Germany) and focused on a camera (Orca Flash v4.0, Hamamatsu Photonics K.K., Japan) with a tube lens (Thorlabs, USA). For further details refer to ^27^.

The piezo, galvanometric mirrors and the triggering of the camera was controlled by Sashimi^65^ a custom-written python software developed in the lab. The lightsheets and the collection objective were synchronously oscillating along the vertical axis with a frequency of 2.0 Hz, covering in depth around 250 µm. Frames were acquired at equally spaced intervals along the volume with a spacing of 9-10 µm.

#### Embedded (tethered) preparation

For lightsheet experiments 6-7 dpf fish are placed in 2.2% low-melting point agarose (Thermofisher) in a chamber optimized for our lightsheet microscope.

The chamber is filled with filtered fish water and agarose is removed along the optic path of the lateral and frontal laser beams (to prevent scattering), around the tail of the animal, to enable movements of the tail and around the mouth and nose. After embedding, fish are left recovering overnight before the imaging session. Before starting the imaging, light tapping on the side of the chamber is used to select the most active fish for the experiment.

The tail of the fish was tracked using an infrared source (RS Components) illuminating the larva from above. A camera (Ximea) was focused on the fish from below and through the transparent bottom of the lightsheet chamber and acquired frames at 400 Hz. Tail movements were tracked online using Stytra^66^.

### Data analysis and Statistics

All analysis were performed using Python 3.8 and relevant Python libraries for scientific computing, like numpy^67^, scipy^68^ and scikit-learn^69^. Dataframes and dataframe manipulations were performed with pandas library^70^.

The figures were produced using matplotlib^71^. For statistical analysis, unless otherwise stated, we used the non-parametric Mann-Whitney U test for unpaired comparisons (*mannwhitneyu* from scipy).

n.s.: not significant, p-value > 0.05

*: p-value 0.05-0.01

**: p-value 0.01-0.001

***: p-value < 0.001

### Behavior

#### Tracking and general preprocessing

For both WT freely swimming and head-restrained experiments the relevant parameters were tracked online using Stytra^66^. For each experiment metadata with all the relevant information were saved in the same folder of the behavior and the stimulus file. Most of behavioral preprocessing (like extraction of swim events) was performed using the python package Bouter^72^. Briefly, swim events were segmented putting a threshold on the velocity of the centroid. We, then, calculated the angle turned as the difference in heading between the beginning and the end of a swim.

For experiments with nacre (mitfa−/−) transparent fish we recorded an .mp4 video, extracted fish centroid with Deeplabcut 2.0^73, 74^, and then preprocessed with custom-made scripts. In particular, we segmented swim events using velocity of the centroid (1mm/s for swim onset) and extracted angle turned by fitting a linear vector using xy trajectories before and after a bout and measuring the signed angle between them.

For head-restrained experiments, we computed the standard deviation of the tail angle trace in a rolling window of 50 ms. Then, using a threshold of 0.1, we found swim onset. The sum of the tail angle during the first 70 ms of a swim has been shown to approximate well the angle turned by a freely swimming fish^29,75^and was therefore used to define angle turned during swims.

We set a threshold of ± 0.52 radians (30 degrees) and we respectively classified swim events (both for freely-swimming and head-restrained experiments) as right, left and forward swims. By convention, negative sign of angle turned is toward the left.

#### Extraction of relevant behavioral parameters for large arena experiment

We flipped horizontally the trajectories when the warm side was on the right. Therefore, by convention, the unpleasant stimulus is always on the left. To express the coordinates of the fish in terms of the temperature experienced we quadratically interpolated temperature calibrations taken every 2 cm in the arena before and after each batch of experiments.

Since all experimental conditions except for the WT preferred a higher temperature, we decided to express the currently experienced temperature as a function of the absolute distance from the fish setpoint. This setpoint was separately computed for WT, chemogenetic experiments and 2-photon ablation experiments. As shown by Supplementary figure 5 b and 6 c, no difference in setpoint was found between experimental and control groups, indicating that this shift in setpoint is independent from the region ablated.

The point of the analysis was to see whether fish behavior was modulated by sensory history and previous motor choices. To this end, we computed, for each bout, the following parameters:

- Angle turned during the swim event (see previous paragraph).
- Experienced temperature before and after swim
- Difference in temperature brought by the previous swim (swim_-1_), if the interswim interval was ≤2 seconds. In the work, we refer to this variable as sensory context.
- Difference in temperature brought by the second-to-last swim (swim_-2_), if the interbout interval between bout_-1_ and bout_-2_ was ≤2 seconds. In the work, we refer to this variable as past sensory context.

Sensory context and past sensory context were further classified in WCxt (worsening context), ICxt (improving context) and NCxt (no context). For such classification, we used a conservative threshold of 0.13°C, which we posit to be the thermal sensitivity in the temporal domain^76, 77^.

Fish that swam less than 1/3 Hz during an experiment were excluded from further analysis.

#### Increase in turn fraction index

To assess whether a fish would be more prone to turn when it experienced a WCxt we took all the swim events happening at least 1 °C away from setpoint and we looked at the fraction of turns happening in a WCxt and in an ICxt. Then, we took the difference between the two contexts.

In this analysis, we did not use the swim events in a NCxt. However, given the shape of our arena and gradient, it was common for the fish to use more than one movement to reverse a WCxt in an ICxt or vice versa (Figure 2g), therefore going through movements during which the temperature did not change (Figure 2i). Despite this, fish still proved efficient homeostatic navigation. To account for this, we repeated the previous analysis, but this time, we selected bouts when sensory context was NCxt but past sensory context was either WCxt or ICxt.

#### Motor correlation index

In order to investigate if fish strengthen directional correlation during WCxt we selected all swims during such sensory context and further selected swims that were preceded by a turn (either left or right). Finally, we computed the theta swim normalized by the direction of the previous turn (we flipped the sign of current swim if previous turn was toward the left), such that if the sign is positive it means that fish moved ipsilateral respect to previous direction, contralateral otherwise.

#### U-maneuvers

In order to extract U-maneuvers, we selected sequences of six consecutive swim and computed the absolute amount of reorientation. If this number exceeded 100°, we considered that sequence a U-maneuver. We further selected U-maneuvers where the first swim was in a WCxt. Finally, we averaged the ΔTemperature experienced during each swim in the sequence.

#### Increase in turn fraction and motor correlation in open-loop head-restrained and freely swimming (small arena)

In these experiments, we tested whether the direction of change (WCxt vs ICxt) was enough to modulate the reorientation probability and directional correlation even if the fish had no control over the experienced temperature change.

For data analysis, we considered two main parts: when the temperature was moving away from 24 °C (24 °C to 28 °C = WCxt) and when it was moving towards 24 °C (28 °C to 24 °C = ICxt). We excluded the step at 30 °C from the analysis. We next computed the increase in turn fraction and motor correlation in the same way as we did for experiment 1 (see “Increase in turn fraction index” and “Motor correlation index”).

#### Coefficient of dispersion

To test whether ablated fish were less able to stay around their setpoint we calculate the coefficient of dispersion:

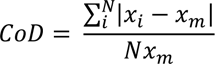

where x_i_ is the preferred temperature of each fish, x_m_ is the median preferred temperature of the population and N is the total number of fish.

### Lightsheet data analysis

#### Lightsheet imaging data preprocessing

Data were saved in hdf5 format. To preprocess functional imaging data we used the package fimpy (https://github.com/portugueslab/fimpy). For alignment (function *align_volumes_with_filtering*) and ROI extraction (function *correlation_map* and *grow_rois*), we used similar pipeline used in ^27^ (see “Whole-brain functional imaging data analysis”). Since planes were spaced by ∼10 µm, ROIs detected across planes were not merged. Once the ROIs were identified, we generated an ID ROI stack: a matrix with the same shape of the lightsheet data where to each pixel was assigned a scalar value corresponding the ID of a single ROI. After ROI detection we removed spurious ROIs detected outside the brain by manually drawing a mask. We further removed all ROIs detected in the olfactory epithelium since it was imaged only in a small fraction of fish and it was prone to many movement artifacts. Alignment was performed independently for both temperature and salt session while ROI extraction only for temperature. After alignment, we additionally computed an anatomical stack by averaging fluorescence throughout the experiment for both sessions (anatomical stack).

#### ROI segmentation across sessions

Registration across sessions was performed by finding an affine transformation matrix semi-manually. This was done by finding ∼15 corresponding single isolated neuronal nuclei in the temperature (used as reference) and the salt session anatomies and computing a least-squares fit of a 12-parameter affine transform matrix. Once found, we applied the inverse transformation (from the temperature to the salt session space) to the ID ROI stack computed in the temperature session in order to extract fluorescence activity from the same neuron in the salt session.

#### Anatomical Registration and region segmentation

Image registration was performed using the free Computational Morphometry Toolkit (CMTK - http://www.nitrc.org/projects/cmtk)^78^. First, we choose one of the anatomical stacks as the initial reference brain, and non-affine volume transformations were computed to align each fish’s anatomical stack to this reference stack using the affine and warp functions^28^. After this step, we averaged all the aligned anatomical stack from all fish in order to obtain an intermediate reference stack. We finally repeated the fish-wise alignment on the intermediate reference stack. These transformations were then used to transform individual ROIs from each fish into the frame of reference of the final reference brain, allowing us to compare the anatomical location of ROIs from different fish. Finally, we built an internal atlas by manually defining anatomical regions using clear anatomical landmarks and by looking at the Max Planck Zebrafish Brain Atlas (https://mapzebrain.org)^79^.

#### Describing temperature sensory responses

Raw traces were first smoothed with a median filter of size 1.5s (function *medfilt*). Then, we cropped all traces from the beginning to the end of each trial (single + double pulse), z-score each of them, and performed an average response per each ROI (Trial trigger average or TTA). In addition, we computed the average correlation (reliability index) of the responses across all individual trials^80^. With the reliability index, we were able to quantify the responsiveness of each ROI to the temperature stimulation independently from each specific response profile. We classified each ROIs as “reliable” if the aforementioned index was higher than 0.3 and further analyzed only this subset of ROIs. This procedure allowed us to keep only neurons reliable across trials. In order to further screen for neuronal responses that were also reliable across animals, we cross-correlated (Pearson correlation) each TTA with all the TTAs of all the other fish and kept only neurons with a coefficient higher than 0.65 with at least one TTA of all the rest of the fish. At the end, we pooled together all the TTAs that passed these screenings. We then performed PCA and projected each TTA on the first two PCs (cumulative explained variance 65%) so that each point in the principal component space was a TTA. Finally, we performed k-means clustering and we set the number of clusters (k parameter) to 3. This number, was chosen by looking at the Davies Bouldin score (*davies_bouldin_score*), lower values of this score indicate better clustering. Clustering quality was also checked by repeating the procedure with different centroids initializations. Regional percentages of different clusters were computed using our internal atlas.

#### Visualization of whole brain maps

In order to generate whole brain maps displayed in the current work we used either a scatter plot or a density map. For the scatter plot (Figure 3 j), color of each dot (representing a ROI) was decided based on cluster identity while transparency was proportional to the local density within a sphere of 30 µm radius. For the density maps (Figure 4 a, f and Supplementary figure 4 e, g), we initialized a zero array with the same size of our internal reference brain (see “Anatomical Registration and region segmentation”). We then increased by one (+1) pixel values corresponding to a spherical region of 15 µm around each selected ROI. For the visualization, we used a sum projection along the three main anatomical axis.

#### Multimodal Neurons

In order to find multimodal neurons, we first selected all ROIs that reliably responded during both temperature and salt session (threshold: 0.3). Anatomical distribution of these ROIs is shown in Supplementary figure 4 g. We then cross-correlated (Spearman correlation) the TTA of the two sessions and, for each ROI, generated a null-distribution by repeating the cross-correlation with circular-shuffled versions of the TTAs. Using the real correlation and the null-distribution we computed a p-value for each ROI. We considered a ROI multimodal if such correlation was higher than 0.3 and the p-value lower than 0.025.

We, then, applied a spatial constrain on the generated density maps shown in Figure 4 f and Supplementary figure 4 g such that we show only regions with an overlap of at least 5 and 10 ROIs respectively.

#### Swim triggered analysis

The analysis aimed to identify neurons whose fluorescence increased after a swim event. First, we selected swim events temporally space 2 seconds from the precedent and 10 seconds from the next. Then we split swims according to the angle turned in order to classify them as left, right or forward swims. Threshold used for classification was the same used for other experiments (±30 degrees). For each fish, number of bout per each category was at least 3. We then cropped each ROI around swim onset (from - 2 to +10s from bout onset). For each ROI, we computed 3 reliability indexes, one for each swim category. We used the 90^th^ percentile of the distribution created by taking the maximum value among the three reliability indexes of all ROIs in order to find a global threshold, within fish, used to define if a ROI was motor responding and specific to one of the three swim categories. In particular, a ROI in order to be selected needed to have one of the three reliability indexes higher than the global threshold and the other two lower than that one so that each neurons was univocally classified as either left, right or forward tuned.

For the map shown in Figure 4 a we then pooled all the left- and right-swim tuned ROIs coming from all fish and computed the density map as described in “Visualization of whole brain maps”. The percentage of motor ROIs was computed using our internal atlas.

Analysis shown in Supplementary figure 4 d aimed to identify different calcium dynamics upon turn onset. In order to do that, we pooled together left- and right-turn tuned neurons and flipped the x coordinates of right-tuned ones. Finally, we sorted turn trigger averages according to timing of peak activity and averaged using temporal bins of 0.5-2.5seconds, 2.5-4 seconds, 4-5.5 and 5.5-10.5 seconds (Supplementary figure 4c).

### Neuron Ablations

#### Chemogenetic Ablations

To perform targeted ablation of Hb, we employed Ntr/NFP pharmaco-genetic approach^51, 81^. Animals expressing nitroreductase (Ntr) in a cell population of interest were treated with prodrug Nifurpirinol (NFP). Ntr converts NFP into a cytotoxic DNA cross-linking agent leading to death of cells of interest. Nirfurpirinol is, like Metronidazole (MTZ), another nitroaromatic antibiotic and it has been shown to reliably trigger cell-ablation at concentrations 2000 fold-lower than MTZ^81^. We used the line *Tg(16715:Gal4v16); Tg(UAS:Ntr-mCherry*), which restricts expression in the dorsal Hb (see Supplementary figure 6 a) and we crossed it with TL wild-type fish. Fish were screened with an upright fluorescence dissecting microscope (Leica, M165 FC) for red fluorescence, at 4 dpf.

mCherry-positive *Tg(16715:Gal4v16); Tg(UAS:Ntr-mCherry +/-)* fish were tested at 5dpf in the freely swimming rectangular arena (Genetic control). In the evening, 5dpf mCherry-positive *Tg(16715:Gal4v16); Tg(UAS:Ntr-mCherry +/-) f*ish and mCherry-negative *Tg(16715:Gal4v16); Tg(UAS:Ntr-mCherry -/-)* (Treatment control) were taken from their Petri dish minimizing the amount of water transferred and placed into a new 9 cm Petri dish filled with 40 ml of 0.2% DMSO (Dimethyl sulfoxide) to increase tissue permeability, 5 µM Nifurpirinol (diluted 1:500 from stock 2.5 mM) and fish water. Solution preparation and fish transfer happened in darkness. Larvae were then placed in the incubator at 28 °C for 16 h in a black box to prevent the inactivation of the drug. The next morning fish were rinsed 3 times to thoroughly remove the NFP and DMSO and were left to recover for one day in fresh fish water.

Positive Hb fish treated with NFP (Experimental group) and negative Hb fish treated with NFP (Treatment control) were then tested at 7 dpf in the freely swimming rectangular arena. After the experiment fish were inspected individually under the fluorescence dissecting microscope to ensure absence of mCherry fluorescence.

The efficiency of Hb ablations was further evaluated by randomly selecting, each time we ran experiments on a different batch of fish, 5-6 mCherry-positive nacre (mitfa−/−) larvae that were imaged under a confocal microscope (Olympus FV1000) before ablations at 5 dpf and then after ablation at 6 dpf and again at 7 dpf (see Supplementary figure 6 b). The rationale of the last step at 7 dpf was to confirm that the process of ablation keeps progressing up to 48 h after treatment with NFP.

#### Laser-mediated cell ablations

At 5 dpf *Tg(elavl3:GCaMP6s+/+) (mitfa−/−)* fish were mounted in 1.5 % agarose, anesthetized with 1x Tricaine (168 mg/L) directly added to fish water and placed under a custom-made 2-photon microscope. For the microscope design and details refer to^48^. To test the involvement of the PoA in thermal navigation we set to bilaterally ablate this structure. The PoA was identified based on its anatomical location and clear anatomical landmarks. Before the experiment, we acquired for each fish a 100 µm stack of the areas we intended to target (see Supplementary figure 5a Pre). Then we used galvo-based scanning to focus 800 nm on a small group of cells at either side of the midline. Even though we target a relatively small group of neurons (5-6 somas) the produced damage extended beyond the targeted area to the nearby cells as also evident by anatomy stacks taken 24 hours after the procedure (and right after the behavioral experiments, Supplementary 5a post). The laser power was 130 mW power measured at the objective back aperture. We developed a protocol where we targeted the group of cells for 200 ms with a 500 ms of interval repeated three times. A Python script automatically controlled both the shutter and exposure time. Neurons were considered successfully ablated when the fluorescence sharply increased and the nuclei looked irregular and fragmented (see Supplementary figure 5a Post). At this point, we took another anatomical image to monitor the localization and extent of the ablation. If the fluorescence of the neurons did not increase or the procedure created air bubbles in the tissue, the fish was not used for subsequent experiments. In control fish, we applied the same protocol used for the PoA to the Optic Tectum. The OT was the only brain region that did not reliably or strongly respond to our temperature stimulus (see Figure 3j). Successfully ablated and control fish were freed from the agarose, returned to a petri dish with fresh fish water and provided with Sera Micron (Sera). Fish were tested 24-48 hours later at 7 dpf. For some fish an additional anatomical stack was acquired after 48 hours to further monitor the scars.

