## Supplementary Figures for "The Preoptic Area and Dorsal Habenula Jointly Support Homeostatic Navigation in Larval Zebrafish"

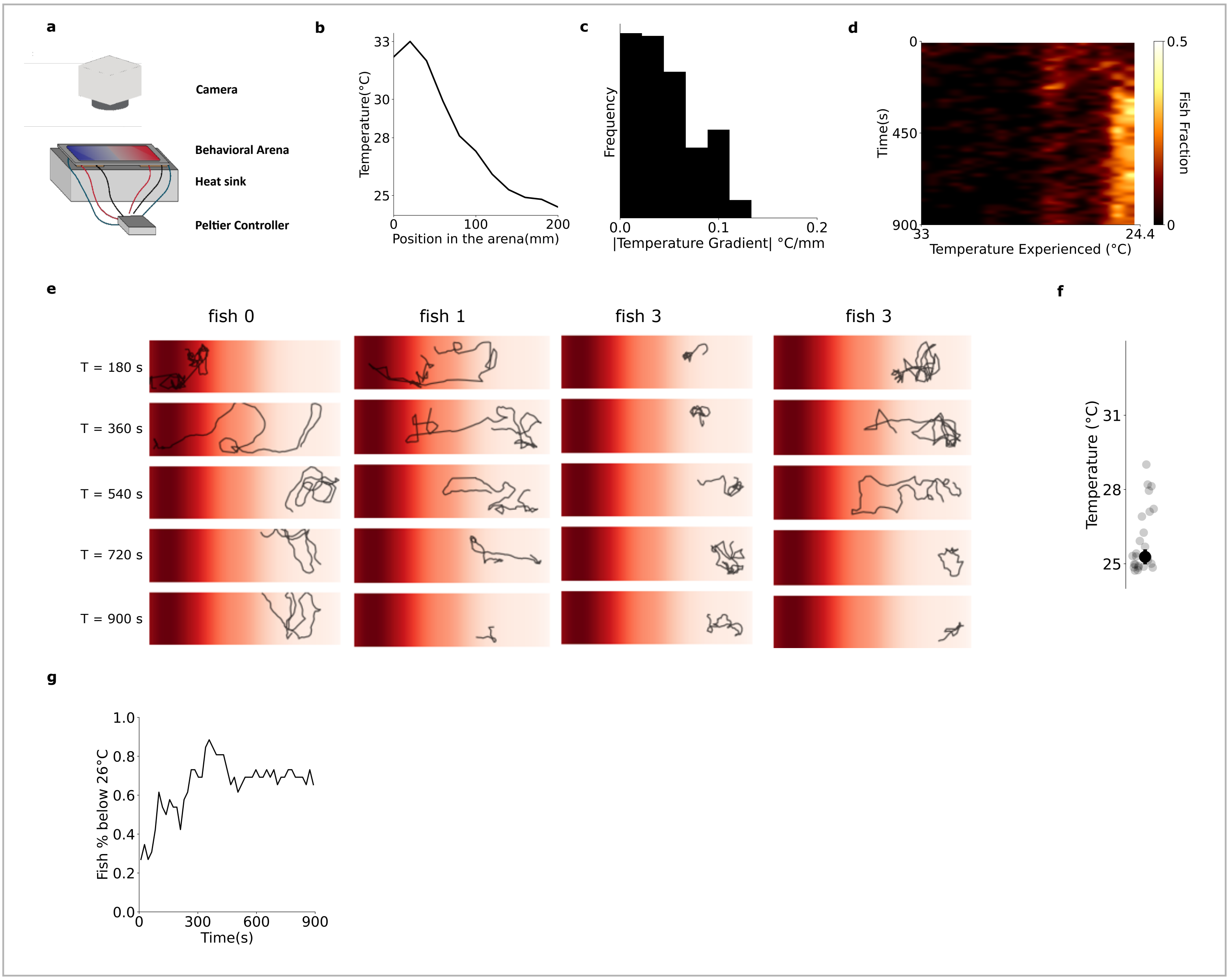


**Supplementary Figure 1**： **a**. Sketch of behavioral setup used for shallow thermal gradient experiments. **b**. Temperature calibration along the arena. **c**. Histogram of temperature gradient along the arena x -axis. **d.** Temperature experienced by fish during different experiment's temporal bins. Color coding represent the fraction of fish . **e**. Example trajectories of 4 selected fish. Trajectories has been split in 3 -minutes bins. **f**. Preferred temperature (setpoint) for WT fish (median±standard error of median). **g**. Fraction of animals placed in the region of the arena where the temperature was below 26 °C.

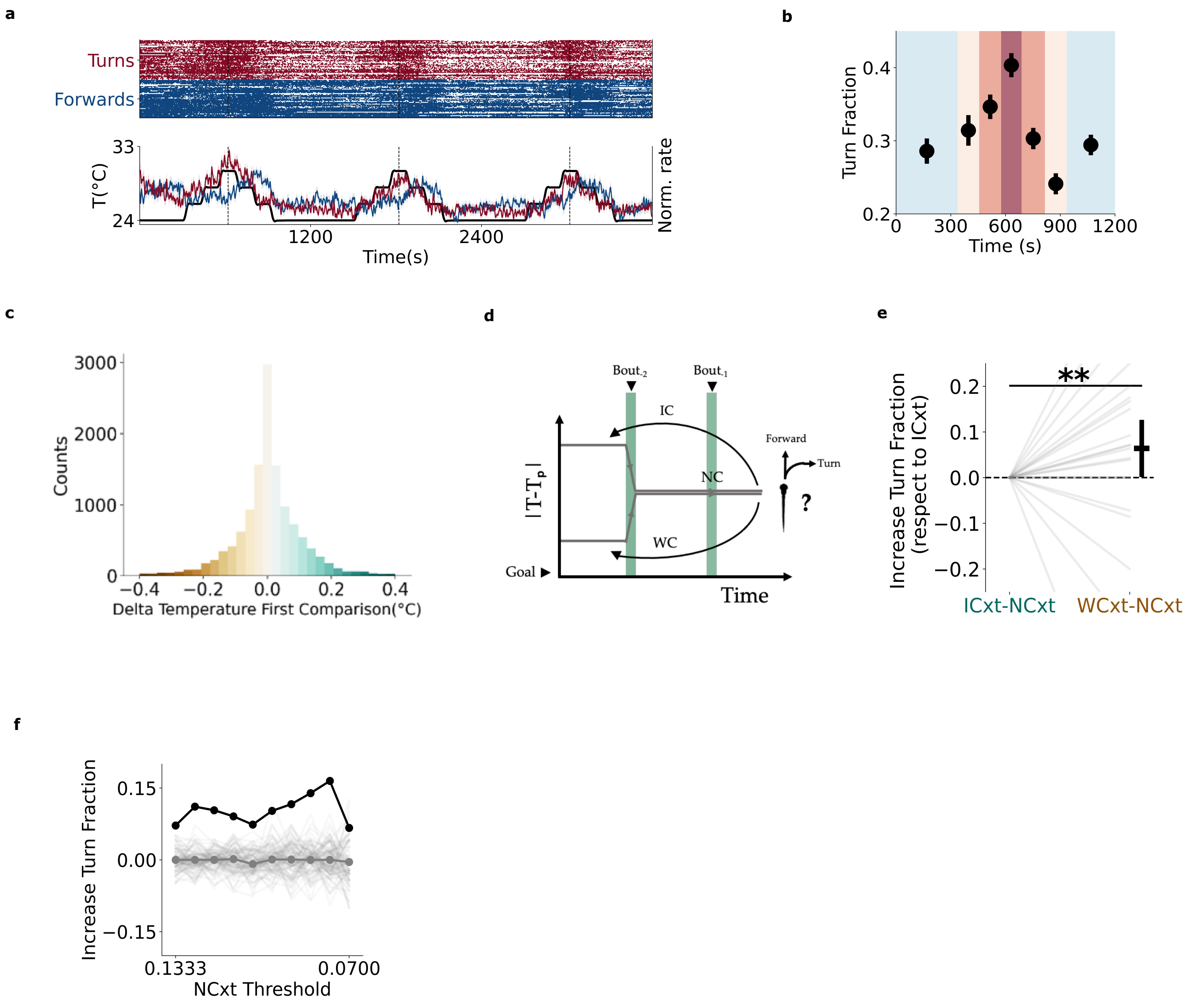


**Supplementary Figure 2**: **a**. Experimental setup used for freely swimming temporal gradient experiments. **b**. Turn fraction according the temperature experienced and sensory context. Color-coding was decided according the temperature delivered. (median ± standard error of median). **c**. Histogram depicting the difference in temperature experienced during between the beginning and end of each swim event. **d**. Sketch depicting the concept of memory of sensory context.
used to define NCxt. **e**. Increase in turn fraction when previous swim did not lead to any directional cues (NCxt) but the previous one was either WCxt or ICxt (median ± standard error of median, Mann-Whitney non parametric test).
**f**. Increase of turn fraction for the WCxt→NCxt condition by changing the threshold.

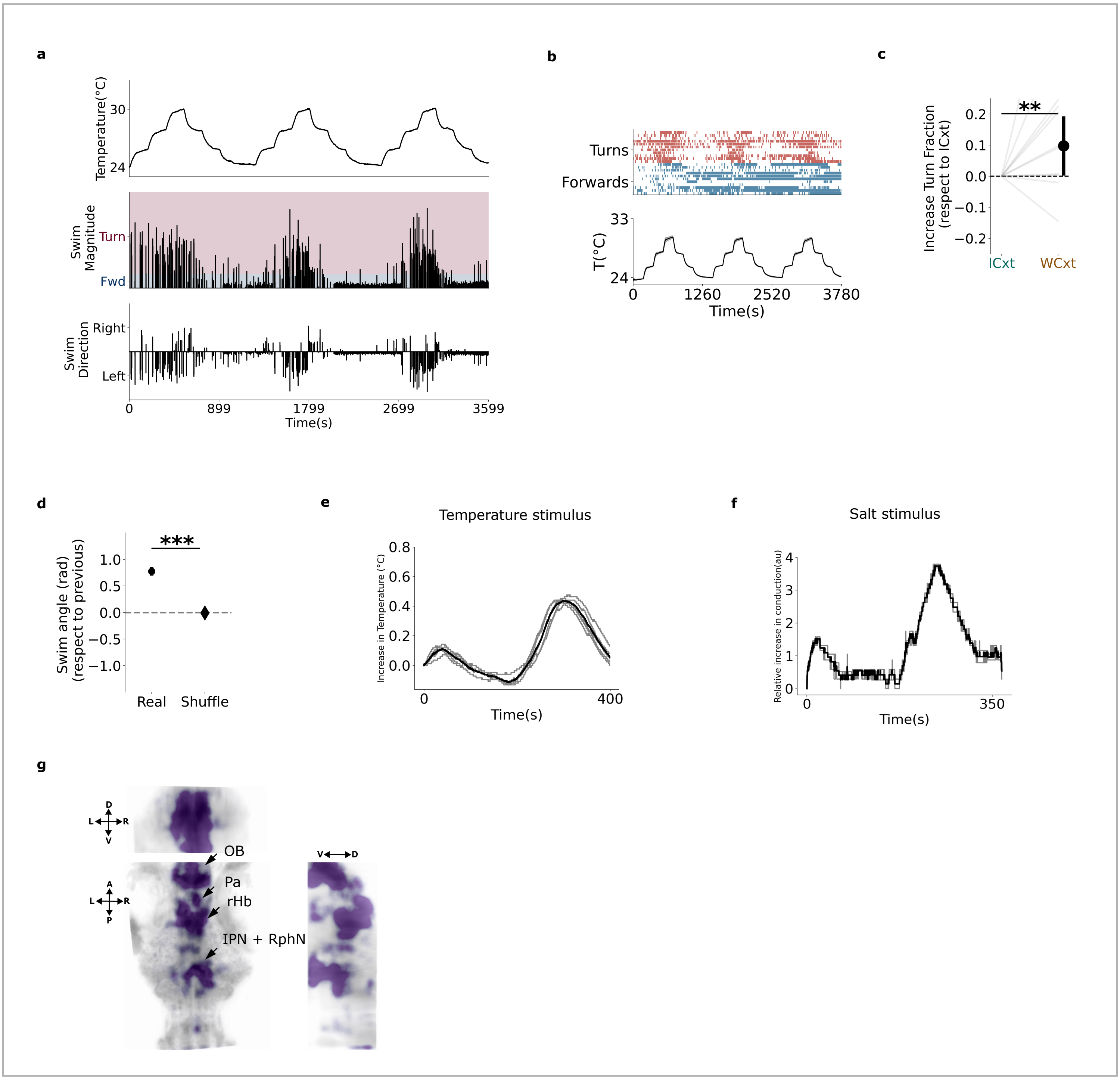


**Supplementary Figure 3: a.** Example fish behavior during head-fixed long temporal gradient experiment. Top: Absolute temperature stimulus delivered to the fish. Middle: Absolute value of reorientation for each swim. Color coding represents: turn (in red, when reorientation is higher than 30 degrees), or forward (in blue, when reorientation is lower than 30 degrees). **IV**: direction of turns, either left or right. **b**.Top: Behavior of fish population (n=11) during head-fixed experiment during long-temporal gradient. Here, we split turn (in red, top) and forward swims (in blue, bottom). Bottom: Temperature stimulus. **c**. Increase in turn fraction depending on sensory context for head-fixed experiments (median ± standard error of median, Mann-Whitney non parametric test). **d**. Turn correlation upon WCxt (mean ± standard error of mean, Mann-Whitney non parametric test). **e**. Temperature stimulus profile for the short protocol used for multimodal experiment**. f.** Salt stimulus profile for the short protocol used for multimodal experiment. **g**. Whole-brain map showing reliable neurons for the long temporal gradient experiment. OB: Olfactory Bulb, Pa: Pallium, rHb: right Habenula, PoA: Preoptic Area, IPN: Interpeduncular Nucleus, RphN: Raphe Nucleus.

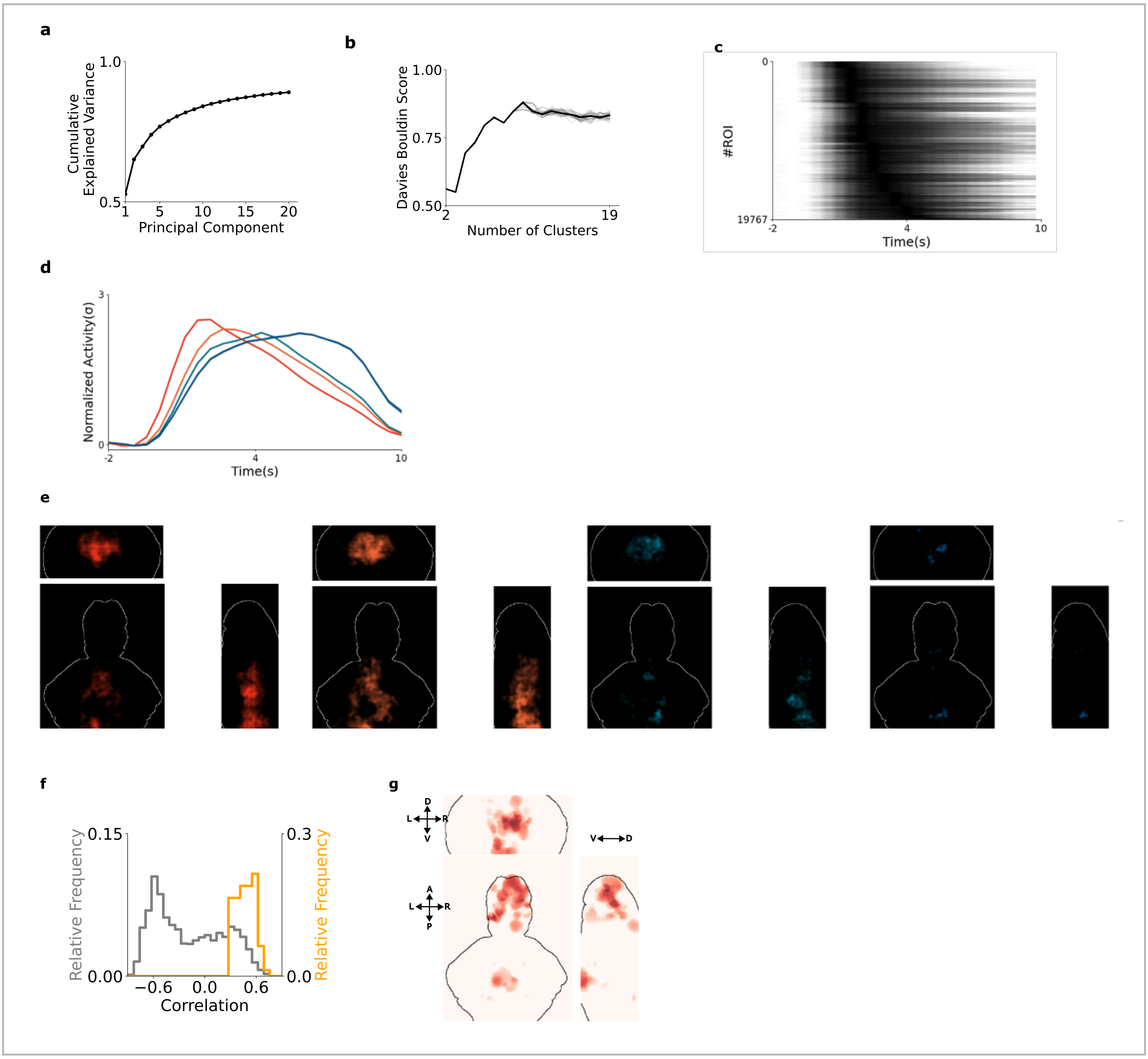


**Supplementary Figure 4** : **a**. Normalized explained variance for the PCA analysis. **b**. Davies-Bouldin score for different number of clusters during k-means clustering analysis. Lower values signify better clusterability with that k. In our dataset, the best k was 3. **c**. Trial turn average for all turn-tuned ROIs. **d**. Mean activity for different lags **e**. Anatomical density distribution for motor ROIs responding with different lags. From left to right: distribution of ROIs with increasing lag. Color coding in the same as in panel **d**. **f**. Distribution of correlation coefficients of ROIs with p -values>0.025 (gray) or lower ( orange). **g**. Anatomical density distribution of neurons reliability responding during both temperature and salt session before the p -value based filtering.

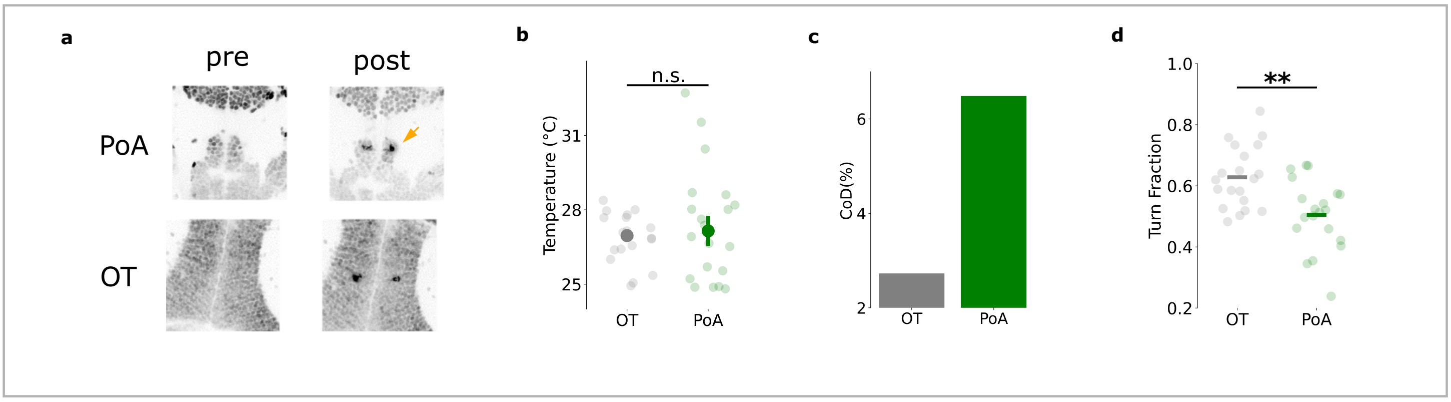


**Supplementary Figure 5** : **a**. Anatomy before (first column) and after (second column) 2 -photon ablation procedure while targeting the PoA (first row) or OT (control, second row). **b**. Coefficient of Dispersion for control group (light gray), ablation group (green). **c**. Preferred temperature for the two experimental groups (median ± standard error of the median, Mann-Whitney non parametric test ). **d**. Median turn fraction computed throughout the entire experiment ( median, Mann-Whitney non parametric test).

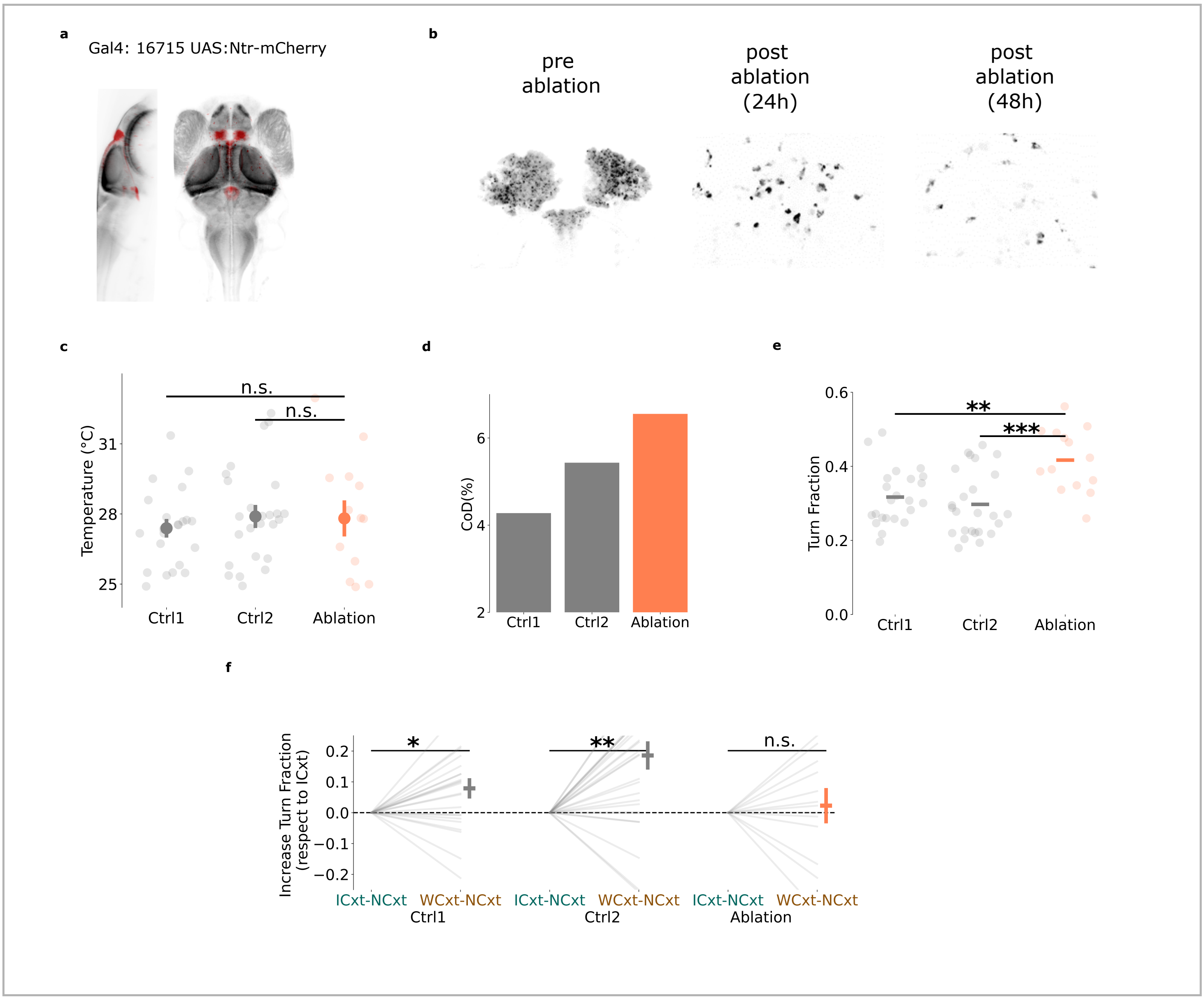


**Supplementary Figure 6**: **a**. Anatomy of a reference brain (gray) and the line Gal4:16715 UAS:Ntr -mCherry (red) used for chemogenetic ablation. **b**. Confocal anatomy of dorsal Habenula acquired before (first column), 24 hours (middle column) and 48 hours (third column) after treatment. **c**. Preferred temperature for the three groups (median ± standard error of the median, Mann-Whitney non parametric test). **d**. Coefficient of Dispersion for control group (grays) and ablation group (coral). **e**. Median turn fraction computed throughout the entire experiment (median, Mann-Whitney non parametric test). **f**. Increased turn fraction when last bout brought a NCxt (median ± standard error of the median, Mann-Whitney non parametric test).
